# A conserved MoxR-vWA chaperone complex drives starvation-induced protein degradation in bacteria

**DOI:** 10.64898/2026.11.14.766676

**Authors:** Hiroyuki Okano, Qiaozhen Ye, Matteo Mori, Gabriel Manzanarez, Zhongge Zhang, Franziska Hackbarth, Lydia R. Chambers, Rohan Balakrishnan, Harish Kannan, Elina Tian, Francisco Mendez-Diaz, Jialu Huo, Eray Enustun, Christina Ludwig, Terence Hwa, Kevin D. Corbett

## Abstract

Bacteria must quickly adapt to changes in their environment, notably including the sudden loss of nutrients. We previously showed that in glucose-starved *E. coli*, widespread starvation-induced protein degradation (SPD) fuels post-starvation proteome remodeling, degrading a wide swath of the proteome to supply the cell with energy and amino acids for synthesis of stress proteins. Here we demonstrate that SPD is mediated by a highly conserved complex of three proteins: the MoxR-type AAA+ ATPase SpdA, the von Willebrand factor A (vWA) domain protein SpdB, and the zinc metallopeptidase SpdC. Structural modeling and *in vitro* reconstitutions show that these proteins form a complex with 6:1:1 stoichiometry that resembles the widespread, but still poorly understood, family of MoxR-vWA chaperone complexes in bacteria and eukaryotes. Measurements of protein degradation in glucose-starved *E. coli* cells show that SpdA, SpdB, and SpdC work together to mediate the degradation of SPD substrates. Instead of directly degrading these substrates, our data suggest that limited structural perturbation and proteolysis “softens” SPD substrates and promotes their degradation by the cellular proteases ClpAP/ClpXP and Lon. Thus, SPD enables extensive proteome remodeling during starvation by expanding the pool of proteins accessible to cellular proteases, rather than by increasing protease abundance. Phylogenetic analysis shows that SpdA, SpdB, and SpdC are highly conserved across bacteria, with notable homologs including the well-characterized *Bacillus subtilis* sporulation factors PrkA (SpdA), YhbH (SpdB), and SpoVR (SpdC).

## INTRODUCTION

All organisms respond to starvation by altering their metabolism. Bacteria frequently encounter nutrient-limiting conditions, inducing a state generically known as stationary phase (1–3) and driving the expression of diverse sets of genes by different bacteria (4–6). Starvation induces some bacteria to form spores, metabolically inactive cells that can resist extended periods of starvation or harsh environmental conditions (7–9). We recently showed that in *E. coli*, carbon starvation induces a nonspecific protein degradation program we term SPD (<u>s</u>tarvation-induced <u>p</u>rotein <u>d</u>egradation) that degrades a broad subset of cytosolic proteins to provide both energy and amino acids for new protein synthesis (10). Under carbon starvation in *E. coli*, SPD requires the general stress factor σ^S^ (RpoS), an RNA polymerase sigma factor that drives the synthesis of several dozen proteins that protect the cell in stationary phase through diverse mechanisms (11). One of these proteins, which we dubbed SpdA (formerly YeaG), is required for SPD. Deletion of *spdA* causes the loss of starvation-induced protein degradation, and strongly reduces the viability of starved cells (10).

*B. subtilis* encodes a homolog of SpdA, PrkA, that plays a crucial but as-yet mechanistically undefined role in sporulation (12). PrkA expression is induced by the sporulation sigma factor σ^E^, which is activated upon nutrient limitation and controls the expression of many sporulation-specific genes (13), and *prkA* deletion causes a strong defect in spore formation (14). Homologs of SpdA/PrkA are found across gram-positive and gram-negative bacteria, suggesting that they carry out a conserved biochemical function that promotes the survival of starving cells, including successful sporulation in certain bacteria.

Based on their amino acid sequence, *E. coli* SpdA and *B. subtilis* PrkA have been previously proposed to function as (i) AAA+ ATPase disaggregases/unfoldases (15); (ii) eukaryotic-like serine/threonine kinases (12, 16, 17); and (iii) standalone proteases (18). The experimental data supporting these proposed biochemical functions are limited and inconsistent, leaving open the question of how exactly SpdA contributes to starvation-induced protein degradation. A cryoelectron microscopy (cryoEM) structure of an SpdA/PrkA homolog from *Thermus thermophilus*, TpkB (*T. thermophilus* protein kinase B), is a homohexameric AAA+ ATPase with homology to MoxR-type or “clade 7” ATPases (15, 19). MoxR ATPases often function as molecular chaperones, together with von Willebrand factor A (vWA) domain proteins. In both *E. coli* and *B. subtilis*, SpdA/PrkA is encoded in an operon with a predicted vWA domain protein: YeaH in *E. coli*, and YhbH in *B. subtilis* (15)*. E. coli* YeaH has been implicated as a functional partner of SpdA (17), and deletion of *B. subtilis yhbH* has been shown to cause defects in sporulation (13, 20). Thus, SpdA/PrkA may be a MoxR-type ATPase that functions alongside a vWA domain protein encoded in the same operon to coordinate SPD during the starvation response.

Here, we combine protein structure predictions, *in vitro* biochemistry, and careful measurements of SPD in carbon-starved *E. coli* to demonstrate that SPD is mediated by a multi-protein complex comprising a MoxR-type ATPase (SpdA), a vWA domain protein (which we dub SpdB), and a predicted zinc metallopeptidase (which we dub SpdC). Structure predictions suggest that this complex resembles other MoxR-vWA chaperone complexes, which play diverse roles in protein homeostasis in both prokaryotes and eukaryotes (19, 21). Structure-guided mutations demonstrate that SPD depends on the predicted active sites of all three SPD complex subunits. Finally, by systematically deleting *E. coli* cellular proteases we find that SPD relies substantially on ClpAP/ClpXP and Lon proteases for its impact on cellular protein turnover, suggesting that the SPD complex does not fully degrade target proteins, but rather primes them for degradation by other cellular proteases. We also find that both the amount of SpdA/B/C proteins and the rate of SPD, which drop over the course of 24 hours after being induced at the onset of starvation, are affected strongly by deletion of the Lon protease. Cells lacking Lon show elevated SPD, suggesting a role for Lon in attenuating SPD during starvation. Overall, our data reveal the architecture and mechanism of a conserved SPD complex that likely destabilizes abundant cytosolic proteins through coupled chaperone-like structural perturbations and limited proteolysis, thereby promoting their degradation by cellular proteases during the starvation response.

## RESULTS

### Identification of a conserved operon controlling stress-induced protein degradation

In *E. coli*, *spdA* (formerly *yeaG)* and *yeaH* comprise an operon (**Figure 1A**) regulated by the stress-responsive sigma factor σ^S^, and prior work has shown that these two proteins function together to respond to different stress conditions (17). The *B. subtilis* homologs of SpdA (PrkA) and YeaH (YhbH) are also encoded in an operon (**Figure 1A**) and are regulated by the alternative sigma factor σ^E^, and both proteins function in sporulation (13, 20). To determine whether SpdA/PrkA and YeaH/YhbH function with additional factors, we pursued two complementary strategies designed to identify genetically linked or physically interacting proteins.

**Figure 1.**
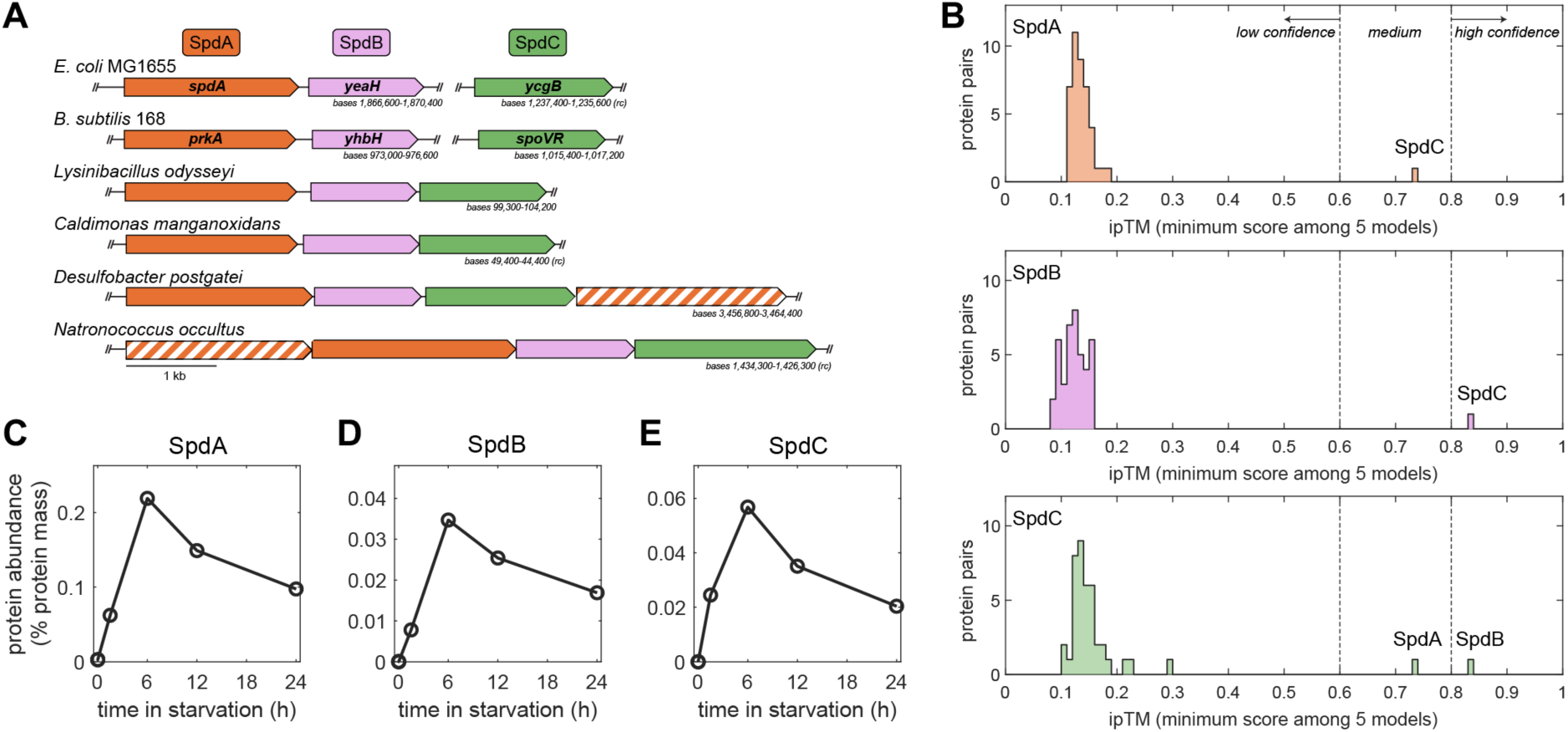
Identification of a conserved bacterial stress-response pathway. (A) Operon schematics of *spdB* homologs in *Escherichia coli* MG1655 (NCBI RefSeq U00096.3), *Bacillus subtilis* 168 (NCBI RefSeq AL009126.3), *Lysinibacillus odysseyi* NBRC 100172 (NCBI RefSeq NZ_BCVX01000011.1), *Caldimonas manganoxidans* ATCC BAA-369 (NCBI RefSeq NZ_KB905947.1), *Desulfobacter postgatei* 2ac9 (NCBI RefSeq NZ_CM001488.1), and *Natronococcus occultus* SP4 (NCBI RefSeq NC_019974.1), highlighting co-occurence of *spdA/yeaG* (orange), *spdB/yeaH* (pink), and *spdC/ycgB* (green) genes across bacteria. See additional examples in **Figure S1A** and **Table S1**. (B) Histograms of AlphaFold 2 interaction screens of *E. coli* SpdA (top), SpdB (middle), and SpdC (bottom) against 42 proteins regulated by σ^S^. Shown is the lowest interface predicted template modeling score (ipTM) out of five independent models (see **Table S2** for full scores). ipTM scores above 0.8 represent high-confidence predictions, between 0.6 and 0.8 represent medium-confidence predictions, and below 0.6 represent low-confidence predictions (83, 84). See **Figure S1B-C** for predicted aligned error (PAE) plots for SpdA-SpdC and SpdB-SpdC predictions. (C-E) Abundance of SpdA (panel C), SpdB (panel D), and SpdC (panel E) after transition to starvation media (data from ref. (10)). The results are expressed as a percentage of total protein mass at the time of measurement. See **Figure S1D-E** for molar ratio calculations.

To identify genetically linked factors, we used the *fast.genomics* server (22) to search for homologs of *E. coli* YeaH across bacterial and archaeal genomes. In a diverse subset of 200 YeaH homologs (out of over 1,600 identified proteins), 176 (88%) encoded homologs of SpdA in the same operon or nearby (**Figure S1A, Table S1**). In 23 cases (including most represented Desulfobacterota and archaeal Halobacteriota), two SpdA homologs were encoded in the same operon. One other gene was strongly associated with *spdA* and *yeaH* in this analysis, being encoded in the same operon in 144 of 200 cases (72%). This gene encodes a homolog of *E. coli* YcgB and *B. subtilis* SpoVR (**Figure 1A, Figure S1A**). In *E. coli* and *Salmonella*, *ycgB* is regulated by σ^S^, suggesting a role in the starvation response (10, 23–26). In *B. subtilis*, *spoVR* is a σ^E^-dependent gene (like *prkA* and *yhbH*) that has been reported to be involved in spore cortex formation (27).

As an alternative, independent method to identify SpdA- and YeaH-associated factors, we performed an AlphaFold 2 interaction screen by predicting the structures of *E. coli* SpdA or YeaH with a set of 42 proteins identified to be in the σ^S^ regulon (**Table S2**) (10). Using the interface predicted template modeling score (ipTM) as a metric, we found that both SpdA and YeaH are confidently predicted to interact with YcgB (**Figure 1B, Figure S1B-C**). A reciprocal screen with YcgB identified only SpdA and YeaH as likely binding partners (**Figure 1B**). Together with our analysis of operonic structure across bacteria, these data suggest that *E. coli* SpdA, YeaH, and YcgB are co-regulated and function together as part of a multi-protein complex. Supporting this idea, mutants of *ycgB* were previously found to have high “co-fitness” with deletions of *yeaG* and *yeaH* in a range of growth conditions with different carbon or nitrogen sources in diverse bacteria (*E. coli*, *Shewanella oneidensis*, and *Pseudomonas fluorescens*) (28).

The above analyses suggest that *E. coli* SpdA, YeaH, and YcgB function together as a complex to mediate SPD, and that this complex is highly conserved across bacteria. We previously developed a method to reproducibly induce carbon starvation in *E. coli* batch cultures, and used this method alongside quantitative mass spectrometry to show that the level of SpdA rises dramatically, then slowly falls, upon starvation (10). Contained in the same dataset are the levels of YeaH and YcgB, which we find follow the same pattern of abundance upon starvation (**Figure 1C-E**). Since our mass spectrometry method is highly quantitative, we can also estimate the relative molar ratio of these three proteins in starving cells at roughly 4:1:1 SpdA:YeaH:YcgB (**Figure S1D-E**). Based on these analyses, we propose that in diverse bacteria, starvation-induced protein degradation is mediated by three proteins that we term SpdA (formerly *E. coli* YeaG/*B. subtilis* PrkA), SpdB (formerly *E. coli* YeaH/*B. subtilis* YhbH), and SpdC (formerly *E. coli* YcgB/*B. subtilis* SpoVR).

### SpdA/B/C are required for protein degradation and survival in carbon starvation

With our system of controlled induction of glucose starvation in *E. coli*, we used radioactivity to measure the total extent of protein degradation (“degradation extent”), defined as the % degradation for proteins synthesized during exponential growth, at various times after the onset of starvation. Briefly, this involves pre-labeling all proteins synthesized during exponential growth by supplying ^14^C-labeled leucine (which is not catabolized by *E. coli*) (29, 30), transferring the culture to glucose-free media with ^12^C-containing leucine, then measuring the amount of ^14^C-labeled leucine that accumulates in the growth media over time (10). Using this method, we found that while wild type cells showed over 20% total protein degradation in the first six hours after onset of glucose starvation, individual deletions of *E. coli spdA*, *spdB*, and *spdC* all showed much lower degradation extent of around 10% after six hours (**Figure 2A, Figure S2**). The rate of degradation was relatively constant in wild type cells over six hours, whereas degradation plateaued after the first 1-2 hours in *spdA*, *spdB*, and *spdC* mutants (**Figure 2A**). This difference is even more stark when visualizing the same data as degradation flux (i.e., rate) over time (**Figure 2B**). In all three mutants, degradation flux shows a simple exponential decrease after the first hour, with a rough half-life of 1 hour, suggesting that degradation targets are becoming exhausted in SPD mutants (approaching ∼12% of the pre-starvation proteome). In comparison, the relative constancy of the degradation flux in wild type cells suggests that the SPD machinery provides an expanded pool of degradation targets, at least through the first six hours after onset of carbon starvation. Consistent with the very similar degradation profiles of *spdA*, *spdB*, and *spdC* mutants, proteomic data obtained from mass spectrometry analysis in these mutants show that their proteomes are very similar to one another during starvation (**Figure 2C-D**).

**Figure 2.**
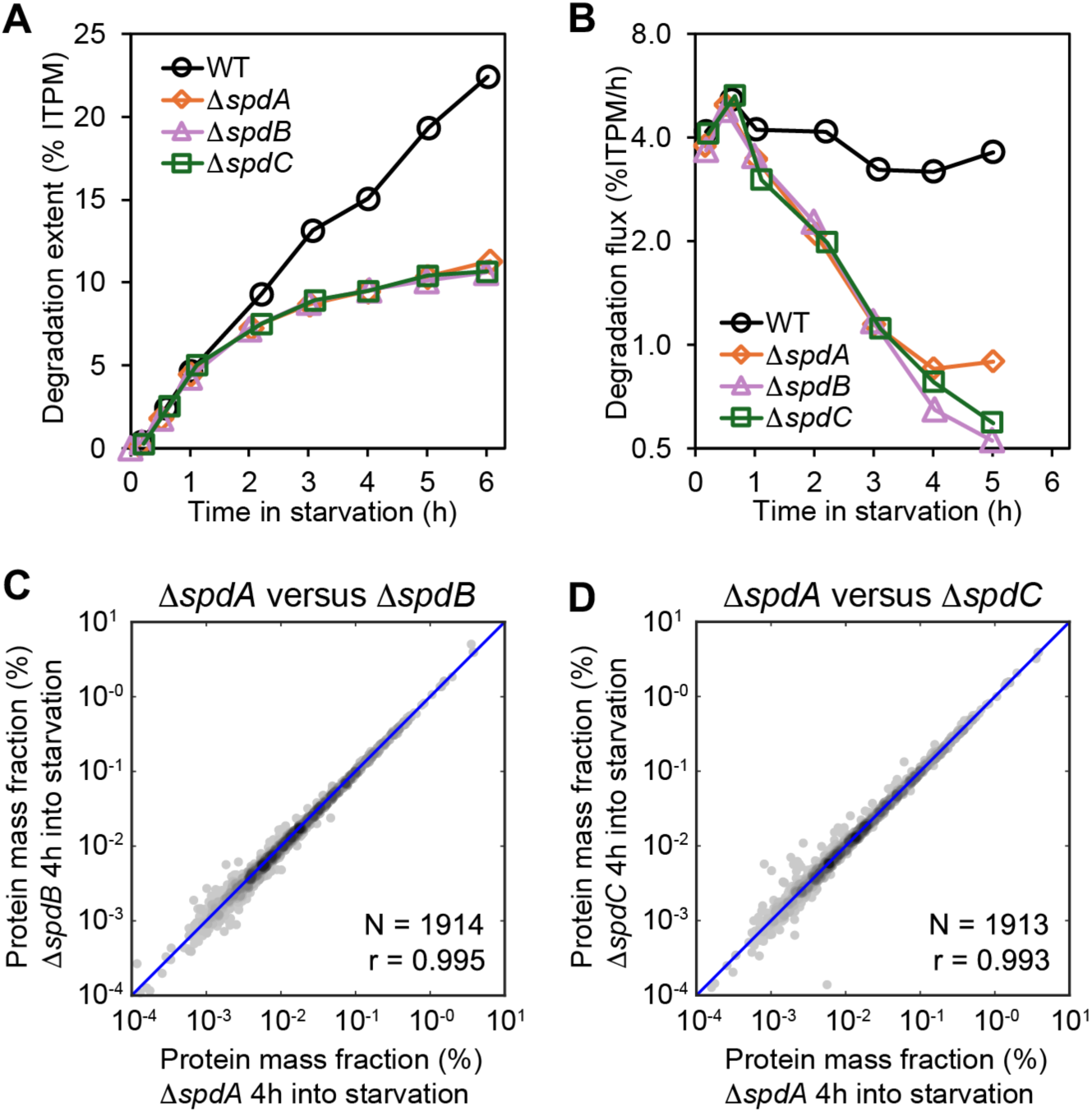
Protein degradation in carbon starvation. (A) Overall protein degradation extent of proteins synthesized during exponential growth for wild type (WT) or indicated mutant *E. coli* cells, as measured by the release of ^14^C-leucine into the medium for cultures grown exponentially with ^14^C-leucine; see **Methods**. ITPM: initial total protein mass (at the onset of glucose starvation). See **Figure S2** for repeatability of degradation extent measurements. (B) Overall protein degradation flux (represented as %ITPM per hour), of wild type (WT) or indicated mutant *E. coli* cells. The data shown are time derivative (1D discrete Laplacian) of the data in panel (A). (C-D) Comparisons of the proteome of Δ*spdB* (panel C) and Δ*spdC* (panel D) mutants with that of Δ*spdA*, 4 h after onset of glucose starvation, when the degradation extent in these mutants is clearly different from wild type cells.

### SpdA is a MoxR-type AAA+ ATPase required for SPD

To learn more about how SpdA, SpdB, and SpdC function together in protein degradation, we used AlphaFold 3 (31) to separately predict the structures of *E. coli* SpdA, SpdB, and SpdC, then used the DALI structure-comparison server (32) to identify structurally related proteins. While SpdA was originally identified as a eukaryotic-like serine/threonine kinase based on limited sequence homology to kinases in its C-terminal domain (12), more recent sequence analyses and biochemical experiments suggest that it is instead a AAA+ family ATPase (14, 15, 18). AAA+ ATPases typically form homohexameric rings involving head-to-tail interactions between their N-terminal “large AAA” domain and the C-terminal “small AAA” domain of a neighboring protomer. The predicted structure of *E. coli* SpdA (**Figure 3A-B**) reveals a large AAA domain spanning residues 81-330, and a small AAA domain spanning residues 331-492. In an AlphaFold 3 prediction of a complex with multiple SpdA protomers, the small AAA domain of each protomer packs against the large AAA domain of its neighboring protomer in a manner similar to other hexameric AAA+ ATPases (**Figure S3A**). In addition to the large and small AAA domains, SpdA is predicted to contain an extended N-terminal region (residues 1-80) that interacts with a small mixed α/β domain insertion in the large AAA domain of a neighboring protomer (residues 130-201). Finally, SpdA is also predicted to contain a short C-terminal α-helical domain (residues 493-644) that does not resemble any known proteins. The predicted structure does not resemble any known kinase family.

**Figure 3.**
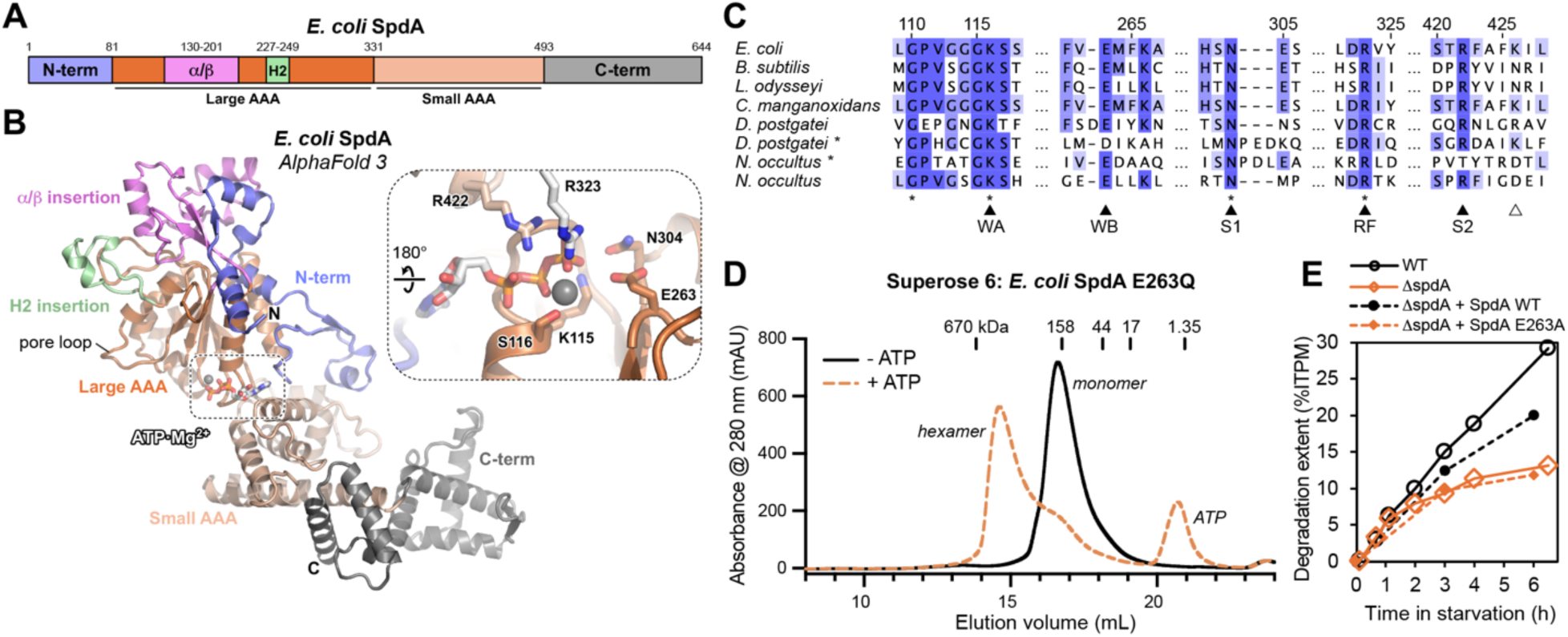
SpdA is a MoxR-type AAA+ ATPase required for SPD. (A) Domain schematic of *E. coli* SpdA, from an AlphaFold 3 structure prediction of three SpdA protomers (**Figure S3**). (B) AlphaFold 3 predicted structure of a single *E. coli* SpdA protomer bound to ATP·Mg^2+^, with domains colored as in panel A and labeled. *Inset:* Close-up view of *E. coli* SpdA ATP·Mg^2+^ binding (view rotated 180° from the overall view), with conserved active-site residues labeled. The arginine finger residue R323 (white) is located on the neighboring protomer. (C) Sequence alignment of ATP-binding motifs in SpdA homologs from Figure 1A (proteins with diagonal shading in Figure 1A are marked with asterisks), with coloring according to conservation in an alignment of 171 SpdA homologs from **Figure S1A** plus *E. coli* SpdA and *B. subtilis* PrkA. Residues conserved in all 173 proteins are marked with asterisks at bottom. Solid arrows indicate ATP-binding residues: WA, Walker A; WB, Walker B; S1, Sensor 1; RF, arginine finger; S2, Sensor 2. Outline arrow indicates residue K426, previously implicated in kinase activity of *E. coli* SpdA (17). Accession numbers for sequences shown are: *E. coli* (UniProt P0ACY3), *B. subtilis* (UniProt P39134), *L. odysseyi* (NCBI WP_036151858.1), *C. manganoxidans* (NCBI WP_019561273.1), *D. postgatei* (NCBI WP_004074833.1), *D. postgatei* * (NCBI WP_004074840.1), *N. occultus* * (NCBI WP_015320726.1), *N. occultus* (NCBI WP_015320727.1). (D) Superose 6 size exclusion chromatography of *E. coli* SpdA Walker B mutant E263Q, in the absence of ATP (black line) or after pre-incubation with 1 mM ATP (brown dashed line). Elution volumes of molecular weight standards are shown at top. Estimated molecular weight in the absence of ATP is 110 kDa (SpdA monomer molecular weight = 74.5 kDa), and in the presence of ATP is 427 kDa (SpdA hexamer molecular weight = 447 kDa). Estimated hydrodynamic radius (Rh) in the absence of ATP is 48.6 Å, close to the 42.2 Å Rh calculated by the HULLRAD server (85) from the AlphaFold 3 predicted structure of a SpdA monomer. Estimated hydrodynamic radius (Rh) in the presence of ATP is 73.4 Å, close to the 74.9 Å Rh calculated by HULLRAD from an AlphaFold 3 predicted structure of a SpdA hexamer. (E) Overall protein degradation extent (ITPM: initial total protein mass) of wild type (WT; black open circles), Δ*spdA* (orange open diamonds), or Δ*spdA* plus a plasmid encoding wild type SpdA (black filled circles) or SpdA E263A (orange filled diamonds). See **Figure S4A-C** for proteomic data showing that the levels of SpdA, SpdB, and SpdC are comparable to those of WT cells when using complementing plasmids. Subtle differences between the proteomes of WT and Δ*spdA* + WT SpdA complemented strain during exponential growth (**Figure S4D-L**) may explain the lack of full recovery of degradation extent in this strain (black filled circles and dotted line).

The predicted structure of *E. coli* SpdA closely matches a recent cryoEM structure of its *T. thermophilus* homolog TpkB (PDB 9XIR; 2.5 Å r.m.s.d. [root mean squared deviation] over 465 C*a* atoms) spanning the extended N-terminal region and large/small AAA domain; the TpkB C-terminus is disordered, precluding a comparison of this domain with *E. coli* SpdA (15). Our *E. coli* SpdA model also closely resembles two recent structures of bacterial MoxR-type ATPases, *Paracoccus denitrificans* NorQ (PDB 9QH8; 3.8 Å r.m.s.d. over 240 C*a* atoms) (33) and *Brucella melitensis* CobS (PDB 9UKG; 3.8 Å r.m.s.d. over 251 C*a* atoms) (34). Both NorQ and CobS form homohexamers and bind one copy of a partner protein possessing a vWA domain (NorD and CobT, respectively) (33, 34). NorQ-NorD functions as a chaperone-like complex, mediating insertion of iron (Fe) into nitric oxide reductase (35). CobS-CobT performs a similar biochemical function, cooperating with the large CobN protein to insert cobalt ions into a tetrapyrrole precursor of vitamin B_12_ (cobalamin) (36–38). The NorQ-NorD and CobS-CobT complexes share a similar architecture, the AAA+ ATPase hexamer wrapping around a “finger” domain extending from the vWA domain of its binding partner. In both NorQ-NorD and CobS-CobT, the vWA finger is engaged by a distinctive β-hairpin insertion into α-helix 2 of the ATPase domain; this insertion is characteristic of so-called “clade 7” AAA+ ATPases (39), and is also present in both the cryoEM structure of *T. thermophilus* TpkB and the AlphaFold 3 model of *E. coli* SpdA (residues 227-249; **Figure 3A-B, Figure S3**). Together, these observations strongly suggest that SpdA is a MoxR-type AAA+ ATPase that functions with a vWA domain protein (likely SpdB; see below).

We expressed and purified *E. coli* SpdA and found that it was monomeric in solution, both in the presence and absence of ATP (not shown). Many AAA+ ATPases only form stable hexamers in the presence of ATP. We generated a Walker B glutamate-to-glutamine mutant (E263Q; **Figure 3C**), which in other AAA+ ATPases prevents ATP hydrolysis and stabilizes the hexamer form (40). SpdA-E263Q was also monomeric, but formed a stable hexamer in the presence of ATP as measured by size exclusion chromatography (**Figure 3D**). Thus, our structure predictions and biochemical data reveal that SpdA is likely a hexameric MoxR-type AAA+ ATPase. Confirming the importance of SpdA-mediated ATP hydrolysis in the *E. coli* starvation response, we found that a Walker B mutant of SpdA was unable to complement a Δ*spdA* mutant in protein degradation extent in glucose-starved cells, despite being expressed at similar levels to the wild type protein (**Figure 3E, Figure S4**).

### SpdB and SpdC form a complex with predicted peptide binding and cleavage domains

We next performed an AlphaFold 3 structure prediction of a 1:1 complex of *E. coli* SpdB and SpdC (**Figure S5A-C**). Consistent with our AlphaFold 2 screen, the two proteins were confidently predicted to bind one another (ipTM = 0.80). In the predicted structure, the N-terminal domain of SpdB interacts with the C-terminal domain of SpdC through two separate short sequences in SpdB that form β-strands and contribute to two different small β-sheets in SpdC (**Figure 4A-B**).

**Figure 4.**
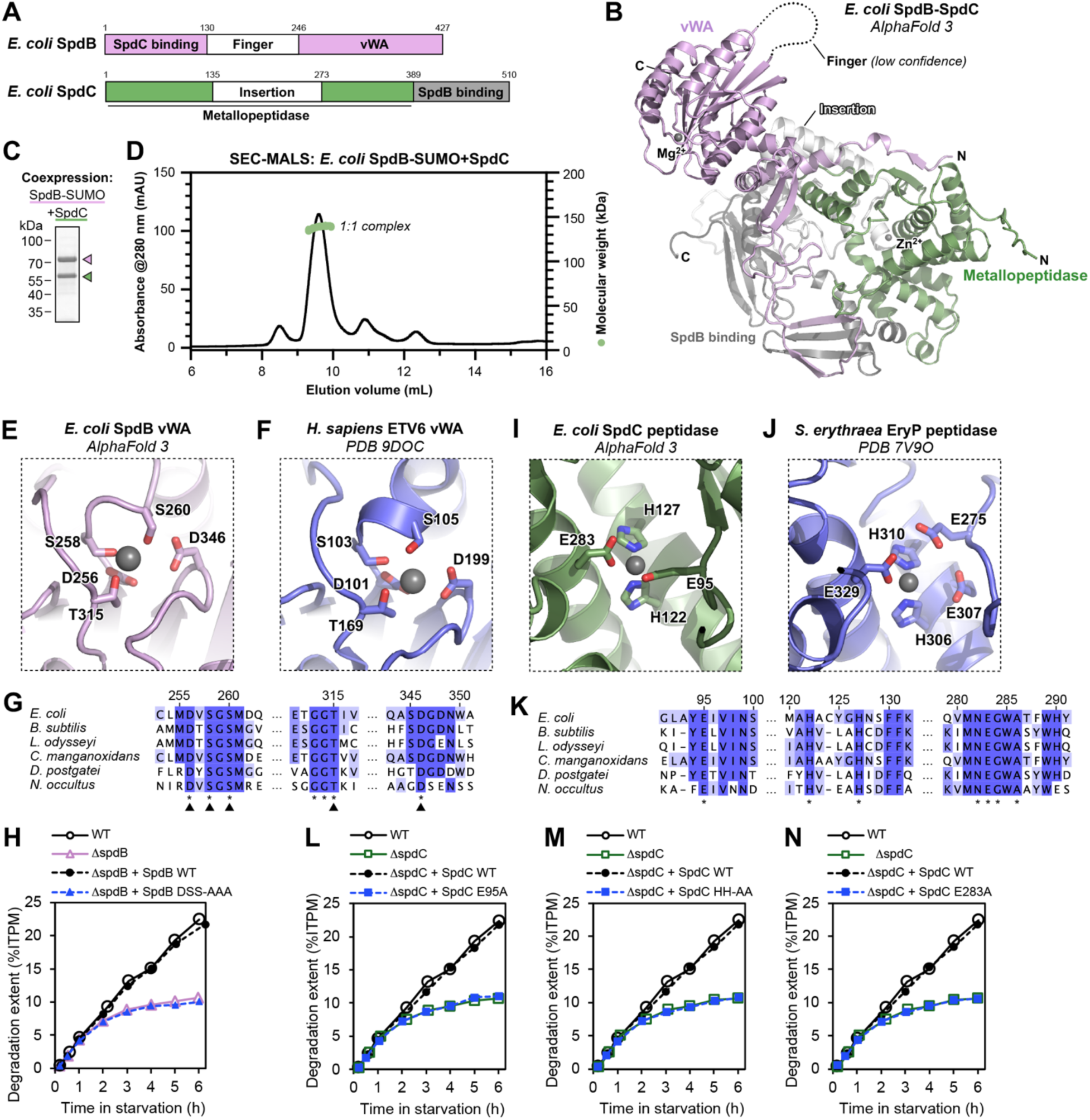
Identification of functional domains in SpdB and SpdC. (A) Domain schematic of *E. coli* SpdB and SpdC, from an AlphaFold 3 structure prediction of a 1:1 complex (**Figure S5**). vWA: von Willebrand factor A domain; Metallopeptidase: M1 zinc metallopeptidase family domain. (B) AlphaFold 3-predicted structure of the *E. coli* SpdB-SpdC complex, with domains of each protein colored as in panel (A) and labeled. The Finger domain of SpdB was predicted with low confidence and is not shown. (C) Coomassie-stained SDS-PAGE gel of purified *E. coli* SpdB-SUMO (pink arrow) and untagged SpdC (green arrow) after coexpression in *E. coli*. (D) Size exclusion chromatography coupled to multi-angle light scattering (SEC-MALS) analysis of a complex of *E. coli* SpdB-SUMO and untagged SpdC. The measured molecular weight (138 kDa) most closely matches that of a 1:1 complex of the two proteins (120 kDa). (E) Closeup of the Mg^2+^ binding site of the predicted vWA domain of *E. coli* SpdB, with conserved Mg^2+^ binding residues shown as sticks and labeled. (F) Closeup of the Mg^2+^ binding site of the vWA domain of *H. sapiens* capillary morphogenesis gene II (PDB ID 9DOC; unpublished), with conserved Mg^2+^ binding residues shown as sticks and labeled. (G) Sequence alignment of Mg^2+^ binding residues in SpdB homologs from Figure 1A, with coloring according to conservation in an alignment of 164 SpdB homologs from **Figure S1A** plus *E. coli* SpdB and *B. subtilis* SpdB. Residues conserved in all 166 proteins are marked with asterisks at bottom. Solid arrows indicate Mg^2+^ binding residues. Accession numbers for sequences shown are: *E. coli* (UniProt P76235), *B. subtilis* (UniProt P45742), *L. odysseyi* (NCBI WP_036151741.1), *C. manganoxidans* (NCBI WP_019561272.1), *D. postgatei* (NCBI WP_004074838.1), *N. occultus* (NCBI WP_015320725.1). (H) Protein degradation extent measurement showing that the loss of SPD in Δ*spdB* is not complemented by a plasmid encoding SpdB with Asp256, Ser258, and Ser260 all mutated to alanine (SpdB DSS-AAA). (I) Closeup of the Zn^2+^ binding site of the predicted M1 zinc metallopeptidase domain of *E. coli* SpdC, with conserved Zn^2+^ binding residues shown as sticks and labeled. (J) Closeup of the Zn^2+^ binding site of M1 zinc metallopeptidase domain of *S. erythraea* EryP (PDB ID 7V9O) (86), with conserved Zn^2+^ binding residues shown as sticks and labeled. (K) Sequence alignment of Zn^2+^ binding residues in SpdC homologs from Figure 1A, with coloring according to conservation in an alignment of 116 SpdC homologs from **Figure S1A** plus *E. coli* SpdC and *B. subtilis* SpdC. Residues conserved in all 118 proteins are marked with asterisks at bottom. Accession numbers for sequences shown are: *E. coli* (UniProt P29013), *B. subtilis* (UniProt P37875), *L. odysseyi* (NCBI WP_036151744.1), *C. manganoxidans* (NCBI WP_019561271.1), *D. postgatei* (NCBI WP_004074839.1), *N. occultus* (NCBI WP_015320724.1). (L-N) Protein degradation extent measurements showing that the loss of SPD in Δ*spdC* is not complemented by mutants in SpdC’s putative metallopeptidase active site: E95A (panel L), His122A/His127A (panel M), and E283A (panel N).

To test whether SpdB and SpdC form a stable complex, we co-expressed *E. coli* SpdB with a C-terminal SUMO-His_6_ tag alongside untagged SpdC, and purified a homogeneous complex containing both proteins (**Figure 4C**). We analyzed this complex using size exclusion chromatography coupled to multi-angle light scattering (SEC-MALS), and obtained a measured molecular weight for the complex of 138 kDa, consistent with the predicted 120 kDa molecular weight of a 1:1 complex of the two proteins (**Figure 4D**).

We next used the DALI structure-comparison server (32) to identify any conserved domains in SpdB and SpdC. This analysis revealed that the SpdB C-terminal domain strongly resembles a vWA domain (**Figure S5D-E**), including those found in *P. denitrificans* NorD (PDB 9QH8; 3.6 Å r.m.s.d. over 157 C*a* atoms) and *B. melitensis* CobT (PDB 21VT; 2.7 Å r.m.s.d. over 171 C*a* atoms). SpdB’s vWA domain also possesses an extended “finger” domain akin to that found in NorD, CobT, and other MoxR ATPase-associated vWA domain proteins (**Figure S6**). vWA domains canonically bind extended peptides of substrates/binding partners via a “MIDAS” (metal ion-dependent adhesion site) motif consisting of the consensus sequence DxSxS; this motif binds a divalent cation (usually Mg^2+^) that engages negatively-charged aspartate or glutamate residues in the binding partner (41, 42). AlphaFold predictions of *E. coli* SpdB bound to a Mg^2+^ ion closely matched crystal structures of vWA domains bound to divalent cations, with a highly conserved DxSxS motif spanning residues 256-260 (DVSGS), and additional conserved residues Thr315 and Asp346 (**Figure 4E-G**). Confirming the importance of this motif, a plasmid encoding SpdB with Asp256, Ser258, and Ser260 all mutated to alanine (SpdB DSS-AAA) did not complement a Δ*spdB* mutant in protein degradation extent in glucose-starved cells (**Figure 4H**).

We next searched for conserved domains in *E. coli* SpdC using the DALI server. This analysis revealed a predicted M1 zinc metallopeptidase domain in the SpdC N-terminus, with the strongest DALI hits including bacterial and eukaryotic aminopeptidases and related enzymes (**Table S4**). Structural overlays with these proteins reveal that SpdC contains a large insertion within the predicted peptidase domain, but is otherwise similar to other M1 zinc metallopeptidases (**Figure S5F-G**). M1 zinc metallopeptidases possess a highly conserved zinc-binding HExxH-(X_18_)-E motif, as well as a GxMEN “exopeptidase” motif that binds and positions the N-terminus of the target peptide (43, 44). *E. coli* SpdC lacks the first glutamate in the HExxH-(X_18_)-E motif, but contains both histidine residues (His122 and His127, spaced one amino acid further than canonical HExxH motifs) and the second glutamate (Glu283; spaced further from HExxH than typical because of the extended insertion in SpdC) (**Figure 4I-K**). These residues are predicted by AlphaFold 3 to bind a Zn^2+^ ion equivalently to other M1 zinc metallopeptidases (**Figure 4I-K**). SpdC mostly lacks a recognizable GxMEN motif, but possesses a conserved glutamate residue in a structurally-equivalent location (*E. coli* SpdC Glu95) that could substitute for the loss of the first glutamate in the HExxH-(X_18_)-E motif (**Figure 4I-K**). We tested the role of all these residues by measuring protein degradation extent in a Δ*spdC* mutant. While a plasmid encoding wild type SpdC complemented protein degradation extent in this mutant, plasmids encoding three different mutants (E95A, H122A/H127A, and E283A) all failed to complement the mutant (**Figure 4L-N**). Overall, our structure predictions, sequence analysis, and protein complex reconstitutions suggest that SpdB and SpdC form a complex with both peptide-binding (vWA domain) and cleaving (M1 zinc metallopeptidase) domains.

### SpdA-SpdB-SpdC is a MoxR-vWA chaperone complex

Our proteomics data show that *E. coli* SpdA, SpdB, and SpdC are all expressed upon carbon starvation, and their relative abundance in starved cells implies a rough stoichiometry of 4:1:1 (**Figure S1D-E**). Based on this observation, the established 6:1 stoichiometry of MoxR-vWA complexes (33, 34), and our demonstration that SpdB and SpdC form a 1:1 complex, we used AlphaFold 3 to model a 6:1:1 complex of SpdA, SpdB, and SpdC, using proteins from both *E. coli* (**Figure S7**) and *B. subtilis* (**Figure S8**). The AlphaFold models of the putative *E. coli* and *B. subtilis* SPD complex are extremely similar to one another, and show a hexameric ring of SpdA protomers, with the single SpdB protomer’s vWA-domain associated finger domain inserted into the SpdA hexamer pore (**Figure 5A**). The SpdB finger domain is directly contacted by a polar residue in pore loop 1 (Asn218; between β-strand 2 and α-helix 2) and by a hydrophobic residue in the extended α-helix 2 insertion (Ile231; **Figure 5B**). The SpdA pore loop and helix 2 insertions form a helical spiral around the SpdB finger domain, with SpdA Ile231 wrapping around a belt of surface-exposed hydrophobic residues in SpdB (**Figure 5B**). The positioning of the SpdB finger domain in the SpdA hexamer pore positions the SpdB vWA domain in the cavity directly below the SpdA hexamer, with the MIDAS motif exposed. In the model, SpdC is sandwiched between the C-terminal α-helical domains of two neighboring SpdA protomers, positioning SpdC’s predicted peptidase active site toward the SpdB MIDAS motif, within the cavity below the SpdA hexamer.

**Figure 5.**
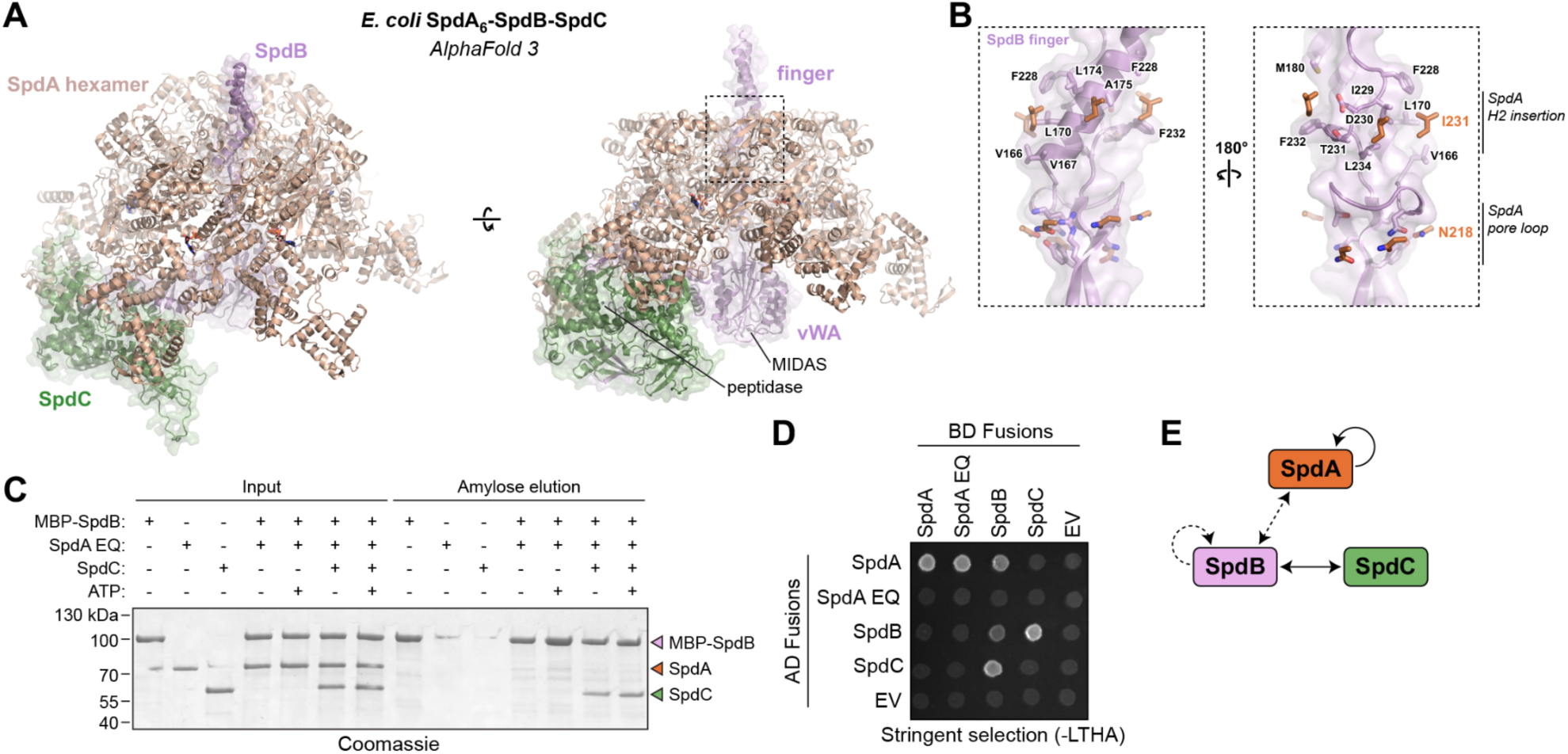
SpdA-SpdB-SpdC is a MoxR-vWA family chaperone complex. (A) Two views of an AlphaFold 3 structure prediction of an *E. coli* SpdA-SpdB-SpdC complex (**Figure S7**). SpdA is colored light brown, SpdB pink, and SpdC green. (B) Two views of the predicted interaction between the SpdB finger (pink) and residues in the SpdA pore loop (N218) and α-helix 2 insertion (H2 insertion; I231). (C) SDS-PAGE analysis of an amylose affinity pulldown assay using His_6_-maltose binding protein (MBP)-SpdB as bait. SpdA EQ: E263Q Walker B mutant. (D) Yeast two-hybrid results for *E. coli* SpdA (WT or E263Q mutant), SpdB, and SpdC. AD: activation domain; BD: DNA-binding domain; EV: empty vector. Shown is growth on stringent-selection plates (-LTHA; missing leucine, tryptophan, histidine, and adenine). See **Figure S12** for growth on non-selection (-LT) and selection (-LTH) plates. (E) Summary of yeast two-hybrid data. SpdA shows self-interaction; SpdB shows self-interaction (dotted line); SpdB and SpdC robustly interact, and SpdA and SpdB interact in one tag combination (dotted line).

The overall architecture of the SpdA-SpdB-SpdC complex closely resembles the recent structures of *P. denitrificans* NorQ-NorD (33) and *B. melitensis* CobS-CobT (34), with the SpdB finger domain inserted into the hexamer pore of the spiral-shaped SpdA hexamer. We also used AlphaFold 3 to model complexes of three other known bacterial MoxR-vWA complexes: *Acidithiobacillus ferrooxidans* CbbQ2-CbbO2, *E. coli* RavA-ViaA, and *Synechocystis* ChlI-ChlD-ChlH. CbbQ2-CbbO2 repairs inhibited active sites of ribulose 1,5-bisphosphate carboxylase/oxygenase (Rubisco) through a poorly-understood chaperone-like mechanism (45–47). The AlphaFold model of this complex (which closely matches a prior low-resolution cryoEM structure (48)) shows a spiral-shaped hexamer of CbbQ2 engaging the finger domain extending from the CbbO2 vWA domain (**Figure S9A**). This model also reveals a previously-undetected M1 zinc metallopeptidase domain in CbbO2, in a further parallel with the SpdA-SpdB-SpdC complex (**Figure S9D-H**). RavA-ViaA performs a chaperone-like activity to maintain the activity of the fumarate reductase electron transport complex (49), and the predicted structure of this complex shows the same overall architecture with a RavA hexamer engaging a single ViaA subunit through its finger domain (**Figure S10**). Finally, *Synechocystis* ChlI-ChlD-ChlH is a metal chelatase complex similar to CobS-CobT-CobN (34, 50). Our predicted structure of this complex includes five copies of the AAA+ ATPase ChlI forming a 5:1 ring with one copy of ChlD, a fusion of an N-terminal AAA+ domain and a C-terminal vWA domain (**Figure S11A-C**). A single copy of ChlH binds the MIDAS motif of ChlD through a negatively-charged glutamate residue (**Figure S11D**). Our AlphaFold model of ChlI-ChlD-ChlH also broadly agrees with a prior low-resolution cryoEM structure of *Rhodobacter capsulatus* BchI and BchD, homologs of *Synechocystis* ChlI and ChlD (51). Overall, these structure predictions reveal that bacterial MoxR-vWA complexes, which play diverse chaperone-like and other roles, have a highly conserved architecture.

To test interactions between *E. coli* SpdA, SpdB, and SpdC, we took two approaches. First, we separately purified all three proteins from *E. coli* and used an affinity pulldown assay with tagged SpdB. We found that SpdB robustly bound SpdC, but did not detectably bind SpdA (**Figure 5C**). We were unable to detect SpdA binding even using the SpdA Walker B mutation (E263Q) in the presence of ATP, which stabilizes the SpdA hexamer. As a second approach, we used yeast two-hybrid assays. In 1:1 interaction tests, we detected robust SpdA self-interactions consistent with hexamer formation, and robust binding between SpdB and SpdC (**Figure 5D-E, Figure S12**). We also detected binding of SpdA and SpdB with one combination of tags (AD-SpdA + BD-SpdB), but not the opposite combination (BD-SpdA + AD-SpdB) (**Figure 5D-E**). Overall, these data suggest that while SpdB and SpdC form a stable 1:1 complex, the association with an SpdA hexamer is likely transient or unstable in the absence of a substrate protein/peptide.

### SpdA-SpdB-SpdC cooperates with cellular proteases to mediate starvation-induced protein degradation

Our biochemical assays, structure predictions, and sequence analyses together suggest that SpdA, SpdB, and SpdC form a complex related to known MoxR-vWA chaperones with an integrated zinc metallopeptidase domain. These data suggest that instead of degrading substrates fully on its own, the SPD complex may use SpdA and SpdB to perturb the structure of a bound substrate through its chaperone-like activity, and couple that to limited proteolysis via SpdC, priming the substrate for further degradation by other cellular proteases.

In carbon-starved cells, all major cellular proteases including ClpAP/ClpXP, Lon, HslUV, and FtsH are largely maintained for at least 24 hours after onset of carbon starvation, regardless of the presence of the SPD complex (**Figure S13**). To test for a role in SPD for each of these proteases, we systematically deleted these genes (with the exception of *ftsH*, which is essential) and measured protein degradation extent during the first six hours of glucose starvation (**Figure 6A-B**). First, we found that deleting *hslV* had no effect on degradation extent (**Figure 6A**). Deleting the AAA+ unfoldases *clpA* and *clpX* each showed a small reduction in degradation, and deleting both *clpA* and *clpX* showed a stronger reduction equivalent to deletion of *clpP*, their shared protease subunit (**Figure 6A**) (52–54). Notably, the reduction in degradation flux in *clpA*/*clpX*/*clpP* mutants was evident within the first hour of carbon starvation, when the SPD complex is not present or contributing to degradation flux (**Figure 1C-E**, **Figure 2A-B**). This observation underscores the point that constitutive protein degradation during the first hour of carbon starvation (10) is implemented by the housekeeping proteases, specifically, the ClpAP/ClpXP system, without assistance from SPD.

**Figure 6.**
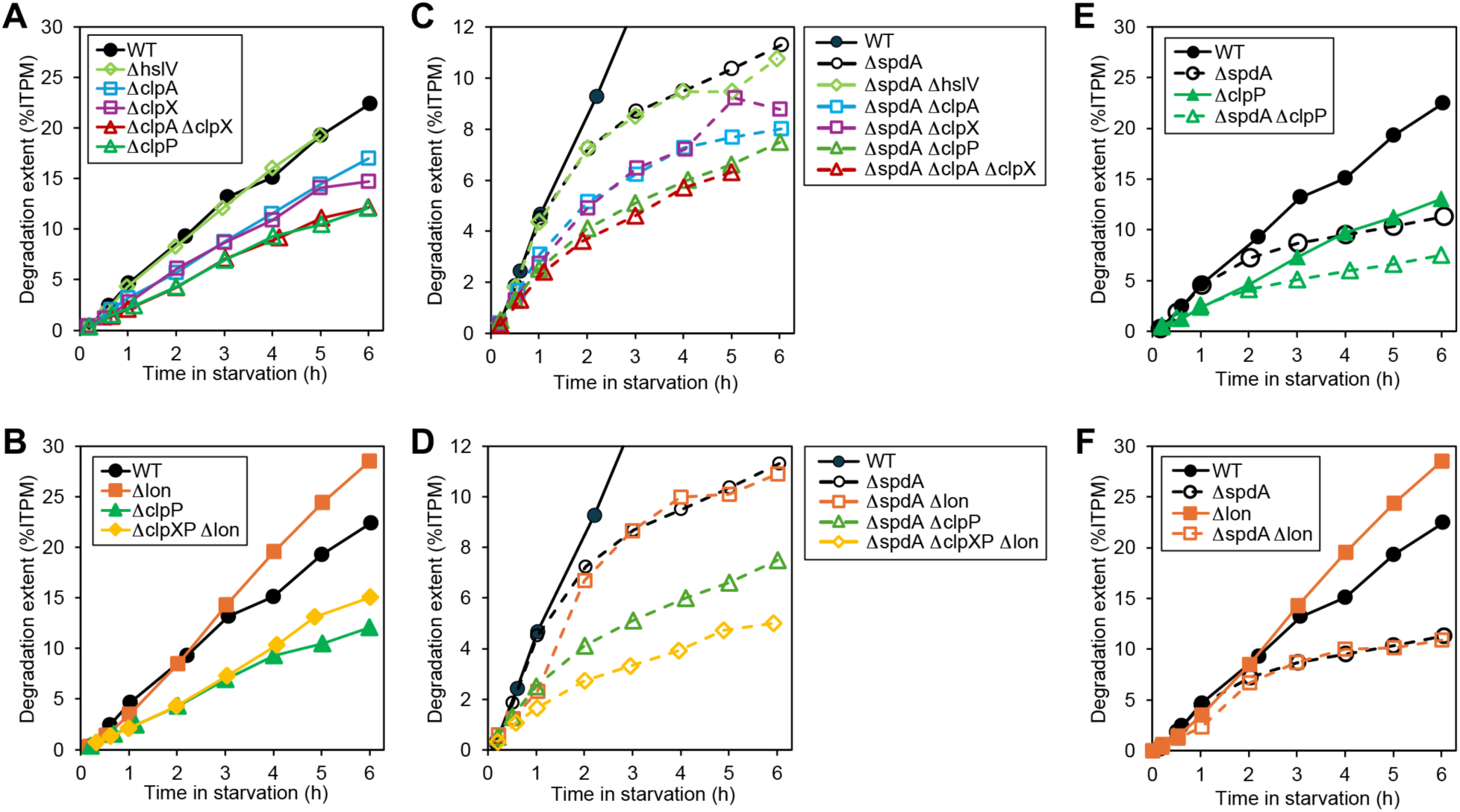
Role of cellular proteases in SpdA-SpdB-SpdC-promoted protein degradation. Degradation extent measurement was performed for combinations of protease deletion mutants and *spdA* deletion mutant. Protease deletion mutants are shown in two groups, those involving Clp proteins (ClpA, ClpX, ClpP) (panel A, C) and those involving Lon (panels B, D). Panels (C) and (D) show the combination with Δ*spdA*. Panel (E) shows a comparison of Δ*clpP* with Δ*clpP* Δ*spdA*, and panel (F) shows a comparison of Δ*lon* with Δ*lon* Δ*spdA*.

We next deleted *lon*, and found that this mutant surprisingly showed an increase in degradation flux over wild type cells after the first two hours (**Figure 6B**). Expressing Lon from a plasmid in Δ*lon* cells reduced degradation extent close to WT cells (**Figure S2D**), confirming a role for Lon in suppressing SPD. One explanation for this effect could be that Lon is involved in the eventual degradation of SPD machinery, whose levels decline after six hours of starvation (**Figure 1C-E**). Alternatively, Lon could degrade σ^S^ to attenuate the starvation response. Consistent with these scenarios, *lon* deletion did not cause an increase in protein degradation during the first hour of glucose starvation (**Figure 6B**), when SPD proteins are low (**Figure 1C-E**). In fact, we observed a slight reduction in degradation flux in Δ*lon* cells in the first hour of carbon starvation compared to wild type, suggesting that Lon, like ClpAP/ClpXP, may contribute to constitutive protein degradation in the initial phase before SPD (**Figure 6B**). Also consistent with the effect of *lon* deletion in enhancing SPD, a Δ*clpXP*-Δ*lon* mutant showed a slight increase in protein degradation extent compared to Δ*clpP* cells (**Figure 6B**). However, we caution that we observed heterogeneous colony size when constructing the Δ*clpXP*-Δ*lon* mutant by phage transduction, suggesting strain instability (not shown). As *hslV* deletion did not show any change in protein degradation compared to wild type (**Figure 6A**), we did not pursue further characterization of Δ*clpXP*-Δ*lon*-Δ*hslV* mutants in which all three non-essential proteases are deleted.

Next, we combined the *spdA* deletion with the above housekeeping protease deletions. As expected, the Δ*hslV*-Δ*spdA* double deletion showed no difference compared to Δ*spdA* (**Figure 6C**). The relative effects of deleting *clpA*, *clpX*, *clpA*+*clpX*, and *clpP* in Δ*spdA* cells are similar to those in wild type cells (**Figure 6C**). Deleting *lon* in Δ*spdA* cells did not affect protein degradation extent compared to Δ*spdA* after two hours (**Figure 6D**), supporting the idea that increased degradation in Δ*lon* arises because of a reduction in Lon-mediated degradation of the SPD machinery itself. Comparing the protein degradation extent in Δ*clpXP* Δ*lon* Δ*spdA* with Δ*clpXP* Δ*spdA* (**Figure 6D**) shows that Lon contributes to protein degradation in early stationary phase in the absence of the SPD and ClpAP/ClpXP systems. Overall, the strain lacking *clpXP*, *lon*, and *spdA* showed the lowest protein degradation extent, with only ∼4% of pre-synthesized proteins degraded after 6 hours of carbon starvation compared to ∼20% in wild type cells. This residual protein degradation may result from the action of the essential protease FtsH which, while mainly targeting membrane proteins, is also known to degrade some cytosolic proteins (55).

A close comparison of protein degradation in specific protease mutants in the presence and absence of SpdA reveals the functional roles of SPD and its interplay with cellular proteases. First, comparing Δ*clpP* with Δ*clpP* Δ*spdA* (**Figure 6E**) reveals the role of Lon – the main remaining protease – in the presence and absence of SPD, and suggests that SPD boosts the proteolytic activity of Lon from ∼7% of proteins to ∼12% (a ∼70% increase) during the first 6 hours of carbon starvation. Similarly, comparing Δ*lon* with Δ*lon* Δ*spdA* (**Figure 6F**) reveals the role of the ClpAP/ClpXP system in the presence and absence of SPD, with their activity boosted from ∼10% of proteins to ∼30% – a ∼three-fold increase – in the presence of SPD. As noted earlier, some of this boost may be due to increased SPD complex levels in the absence of Lon. Overall, these data show that ClpAP/ClpXP and Lon all contribute to the constitutive phase of protein degradation soon after the onset of carbon starvation, and that SPD feeds both proteases in the extended phase of degradation.

## DISCUSSION

Here we reveal the architecture and function of a highly conserved starvation-induced protein degradation (SPD) complex that promotes large-scale protein degradation in starving bacterial cells. The SPD complex comprises three proteins: the MoxR ATPase SpdA, the vWA domain protein SpdB, and the zinc metallopeptidase SpdC. These proteins are highly conserved across bacteria, and are often encoded in a single operon (**Figure 1A, Figure S1A**). In cases where these proteins span multiple operons (as in *E. coli* and *B. subtilis*), they are co-regulated, induced by stress-responsive regulators like σ^S^ in *E. coli* and σ^E^ in *B. subtilis*. In many bacteria, homologs of SPD components are induced upon starvation and contribute to the starvation response and/or sporulation. Reported examples include *Salmonella enterica* (24), *Shewanella oneidensis* (28), *Pseudomonas fluorescens* (28), *Clostridium beijerinckii* (56), *Rhizobium etli* (57), *Sinorhizobium meliloti* (58), and *Myxococcus xanthus* (59). The predatory bacterium *Bdellovibrio bacteriovorus* expresses homologs of SpdA, SpdB, and SpdC along with homologs of ClpX and ClpP in its “attack phase”, which is also triggered by starvation (60, 61).

Our structural modeling and supporting biochemical experiments reveal the SPD complex to be a 6:1:1 complex of SpdA, SpdB, and SpdC, with a hexamer of the MoxR-type ATPase SpdA binding the finger domain of the vWA domain protein SpdB. SpdB forms a constitutive 1:1 complex with the zinc metallopeptidase SpdC, which also interacts directly with the C-terminal domains of SpdA. We show that the ATPase activity of SpdA, the substrate-binding MIDAS motif of SpdB, and the metallopeptidase active site of SpdC are all required for large-scale proteome degradation in carbon-starved *E. coli* cells.

Across diverse bacteria, complexes comprising a MoxR-type ATPase and a vWA domain protein play specific chaperone-like roles to maintain key molecular machinery in the cell. These complexes include CbbQ2-CbbO2 (which we report incorporates a peptidase domain), RavA-ViaA, NorQ-NorD, and CobS-CobT. Recent experimental structures (33, 48) and our structural modeling suggest that these complexes share a common overall architecture, with an ATPase hexamer engaging the finger domain of its associated vWA domain-containing binding partner. While the specific molecular mechanisms of these machines are not well understood, they are all posited to bind their substrates via the vWA domain MIDAS motif, then couple ATP hydrolysis in their ATPase subunits to structural perturbations in the substrate. Based on this model, we propose that the SPD complex binds diverse substrates in the starving proteome via SpdB, perturbs their structure via SpdA ATP hydrolysis, and performs limited proteolysis via SpdC. This process of structural perturbation and proteolysis may “soften” the substrate proteins, thereby promoting their degradation by the cellular proteases ClpAP/ClpXP and Lon (**Figure 7**). Interestingly, *lon* deletion led to an increase in SPD-dependent degradation, suggesting a role for Lon in SPD attenuation, either by reduced expression of SPD proteins (e.g., by degrading σ^S^) or by directly degrading the SPD complex.

**Figure 7.**
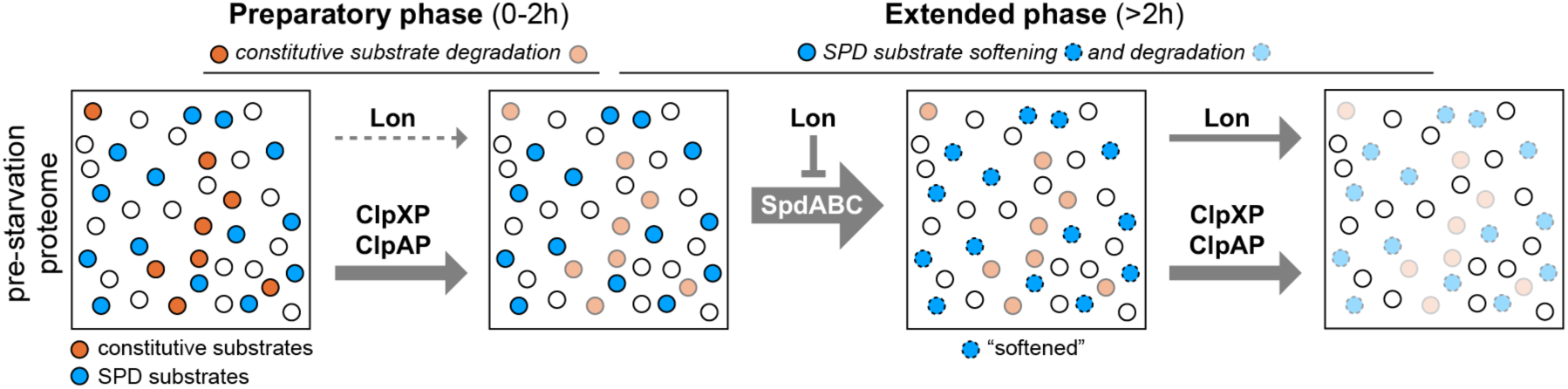
Model for SpdA-SpdB-SpdC in stress-induced protein degradation. In the preparatory phase (0-2 hours after onset of starvation), cellular proteases degrade constitutive substrates (orange circles), with degradation activity dominated by ClpAP/ClpXP. Based on data in Figure 2A-B, constitutive substrates make up ∼12% of the proteome inherited from exponential growth. In the extended phase (more than 2 hours after onset of starvation), the SPD complex softens additional protein substrates (blue circles) through a combination of chaperone-like structural perturbation and limited proteolysis, thereby priming these substrates for further degradation by cellular proteases. Lon suppresses SPD, potentially by suppressing the synthesis of the SPD complex or by degrading this complex during starvation.

An important implication of the mechanism revealed here is that large-scale proteome degradation can be regulated through substrate accessibility rather than protease abundance. By rendering a broad range of cytosolic proteins susceptible to degradation by existing housekeeping proteases, the SPD machinery enables extensive proteome remodeling following nutrient depletion. This strategy may be particularly advantageous during starvation, when the cell must rapidly redirect its limited resources toward the synthesis of proteins needed for survival. The conservation of SPD proteins across bacteria, including their involvement in sporulation in *B. subtilis*, raises the possibility that regulated substrate accessibility represents a more general strategy for proteome remodeling during major physiological transitions. Indeed, sporulation involves an extensive reorganization of the cellular proteome as vegetative cells differentiate into specialized, stress-resistant spores. The involvement of SPD proteins in this process suggests that the same molecular machinery may facilitate proteome remodeling during developmental transitions, extending its physiological role beyond the starvation response.

MoxR-vWA chaperones are widespread across kingdoms (19, 21), including several important but understudied examples from eukaryotes. As one example, Midasin is a large protein (*H. sapiens*: 5596 amino acids) that serves as a ribosome biogenesis factor, mediating the removal of specific assembly factors from ribosome assembly intermediates (62–64). Midasin possesses six tandem MoxR-type AAA domains, an extended central linker domain, and a C-terminal vWA domain. Medium-resolution cryoEM structures of *S. pombe* Midasin (Mdn1) and *S. cerevisiae* Midasin (Rea1) both show the protein’s vWA domain docked against the underside of the AAA+ domain hexamer, tethered by an extended linker that could not be definitively assigned (63, 65). Our AlphaFold model of *H. sapiens* Midasin suggests that this density represents the finger domain associated with the protein’s C-terminal vWA domain, which is engaged by the AAA+ domain hexamer similarly to other MoxR-vWA complexes (**Figure S14**). As another example, Vwa8 is a poorly-understood protein that localizes to the inner mitochondrial membrane and may regulate the activity of electron transport complexes (66). Our AlphaFold modeling shows that Vwa8 has three tandem MoxR-type AAA domains and a C-terminal vWA domain (**Figure S15**). Vwa8 may form a homodimer, or associate with another MoxR-type ATPase, to form the canonical ATPase hexamer assembly engaged with one vWA domain. Both Midasin and Vwa8 are members of a family of AAA+-vWA proteins originally identified over 20 years ago; this family also includes a metal chelatase from *Arabidopsis thaliana* that possesses both AAA+ and vWA domains (42).

A recent comparative genomics analysis in bacteria identified a large set of putative conflict systems (possibly mediating bacteria-bacteria, bacteria-bacteriophage, or bacteria-host conflicts) that include MoxR-type ATPases and vWA-domain proteins linked to diverse putative sensors and effectors (67). This points to MoxR-vWA pairs as conserved conformational signaling modules that can be adapted to a range of contexts.

Our work demonstrates that the bacterial three-protein SPD complex plays critical and highly conserved roles in protein degradation during the starvation response. Many questions remain unanswered, however. One critical question is how SPD targets are recognized. While SPD targets a broad set of bacterial proteins, our prior work showed that metabolic proteins are particularly affected by SPD (10), suggesting that SPD substrates may share a common motif or structural feature recognized by the SPD complex. Alternatively, inactivity of certain metabolic complexes during starvation may prime them for degradation, perhaps due to lower stability of substrate-free apoenzymes versus substrate-bound holoenzymes (68). Another key question is how the SPD complex exerts its pro-degradation effects. Our results suggest that a combination of chaperone-like structural perturbations and limited proteolysis promotes SPD substrates’ degradation by cellular proteases, but the molecular nature of these effects is unknown. On a broader note, the mechanisms of MoxR-vWA complexes as a whole remain mostly mysterious, specifically how these proteins couple ATP hydrolysis to structural perturbations in their substrates to mediate diverse biochemical activities including metal insertion and protein complex assembly and disassembly.

## Methods

### Strains and plasmids

Strains used in this study are described in **Table S5**. The wild type strain is NCM3722 (69, 70); all other strains were derived from this parental strain. Single structural genes spdB (*yeaH), spdC (ycgB), hslV, clpP, clpA, clpX,* and *lon* were individually deleted from NCM3722 by P1 transduction using appropriate kanamycin (Km) resistant mutants from Keio collection as donors. This yielded seven deletion mutant strains HE988, HE997, HE1017, HE1113, HE1114, HE1115, and HE1138, in which *spdB*, *hslV, clpP, clpA, clpX, spdC,* and *lon* were deleted, respectively (**Table S5**). When desired, *E. coli* cells were transformed with plasmids by electroporation or TSB transformation (71).

### Multiple knockout strains involving *spdA* deletion

The following four double mutants were made by P1 transducing the desired mutation to a recipient strain harboring *spdA* deletion (HE845 from ref. (10)) with the Km resistance gene (*kan*) of the latter flipped out from P1 transductant by pCP20 (72): (i) HE1119 (deleted for *clpA* and *spdA* by transferring the *clpA* mutation into the *spdA* single mutant); (ii) HE1120 (deleted for *clpX* and *spdA* by transferring the *clpX* mutation into the *spdA* single mutant); (iii) HE1136 (deleted for *clpP* and *spdA* by transferring the *clpP* mutation into the *spdA* single mutant); (iv) HE1140 (deleted for *lon* and *spdA* by transferring the *lon* mutation into the *spdA* single mutant); and (v) HE1189 (deleted for *hslV* and *spdA* by transferring the *hslV* mutation into the *spdA* single mutant). Strain HE1189 (harboring deletion of *hslV and spdA*) was obtained by transferring the *spdA::kan* allele into the *hslV* mutant HE997. Strain HE1134 (harboring deletion of *clpA, clpX,* and *spdA*) was obtained by flipping out Km resistance from HE1119 and transferring *clpX* mutation into the *clpA spdA* double mutant.

### Multiple protease knockout strains

HE1117 (harboring *clpA/clpX* double deletion) was obtained by transferring the *clpX* mutation into the *clpA* single mutant (HE1113) after flipping out Km resistance of the latter. For triple protease mutation, a strain annotated as *lon*/*clpP*/*hslV* triple mutant was obtained from the Markus Basan Lab (Harvard). PCR and sequencing analyses revealed that this mutant carries the complete *clpP* gene. Thus, this strain is actually a *lon*/*hslV* double mutant (deleted for *lon* and *hslV*) and was therefore not used.

For the *clpP/clpX/lon* mutant, the contiguous chromosomal region carrying *clpP*, *clpX* and *lon* was deleted using the Lambda-Red method (72). The *kan* gene was amplified from pKD4. The purified PCR products were electroporated into NCM3722 cells expressing Lambda-Red proteins encoded by pKD46. Km resistant colonies were confirmed by PCR for the replacement of genes *clpP*, *clpX* and *lon* by *kan*. This yielded strain HE1187, in which *clpP*, *clpX* and *lon* are deleted. Strain HE1186 (harboring *clpP*, *clpX*, *lon*, and *spdA*) was obtained by transferring the ΔΦ(*clpP*:*clpX*:*lon*)::*kan* allele from HE1187 into the *spdA* single mutant with *kan* flipped out.

### *spdA* plasmids

The plasmids pZS-*spdA^+^* and pZS-*spdA1* were constructed as follows. The 343 bp sequence upstream of the 1^st^ codon of *spdA* and the *spdA* coding sequence including the stop codon were tandemly fused and cloned as an XhoI-BamHI fragment on the pZS vector where XbaI of pZS21 (73) was replaced with BamHI. The *spdA1* allele carries a single nucleotide substitution of A788C.

Fusion PCR was used to introduce the *spdA1* mutation. First, two fragments sharing a 43-bp overlapped region in between, p*spdA*-*spdA*’_E263A (containing the 343 bp sequence upstream of the 1^st^ codon of *spdA* and a portion of the *spdA* coding sequence spanning from the 1^st^ codon to the *spdA1* mutation at the 3’ end) and E263A_*’spdA* (containing a portion of the *spdA* coding sequence with the *spdA1* mutation at the 5’ end to the stop codon) were individually amplified from the NCM3722 genomic DNA. These two fragments were then fused together by PCR to obtain the fragment containing the 343 bp sequence upstream of the 1^st^ codon of *spdA*, the *spdA* coding sequence with the *spdA1* mutation, and the stop codon.

### SpdB and SpdC plasmids

The plasmids pZS-*spdB^+^* and pZS-*spdB1* were constructed as follows. The 341 bp sequence upstream of the 1^st^ codon of *spdA*, the 1^st^ 150 bp of the *spdA* coding sequence, the stop codon of *spdA*, the 112 bp intergenic sequence between *spdA* and *spdB* coding sequences, and the *spdB* coding sequence including the stop codon were tandemly fused and cloned as an XhoI-BamHI fragment on the pZS vector where XbaI of pZS21 (73) was replaced with BamHI. The nucleotide substitutions in *spdB1* are A767C, T772G, and T778G.

The plasmids pZS-*spdC^+^*, pZS-*spdC1*, pZS-*spdC2*, pZS-*spdC3* were constructed as follows. The 329 bp sequence upstream of the 1^st^ codon of *spdC* and the *spdC* coding sequence including the stop codon were tandemly fused and cloned as an XhoI-BamHI fragment on the pZS vector where XbaI of pZS21 (73) was replaced with BamHI. The nucleotide substitutions in *spdC1*, *spdC2*, and *spdC3* are A284C, C364G A365C C379G A380C, and A848C, respectively. All plasmid construction was performed by GenScript (GenScript, Piscataway, USA).

### Lon plasmid

The plasmid pZS-*lon* was constructed as follows. The 133 bp sequence upstream of the 1^st^ codon of *lon* and the *lon* coding sequence including the stop codon were tandemly fused and cloned as an XhoI-BamHI fragment on the pZS vector where XbaI of pZS21 (73) was replaced with BamHI.

### Media and growth

Unless otherwise stated, all growth and starvation experiments in this work were performed in MOPS-buffered minimal medium (74) consisting of 40 mM MOPS and 4 mM Tricine (both adjusted to pH 7.4 by NaOH), 10 mM NH_4_Cl, 0.1 mM FeSO_4_, 0.276 mM Na_2_SO_4_, 0.5 μM CaCl_2_, 0.523 mM MgCl_2_, 50 mM NaCl, and micronutrients as described in ref. (75). Glucose was supplemented as a carbon source at the concentration indicated for each experiment.

Growth and starvation were performed at 37°C in water baths shaking at ∼240 rpm. Cell density was measured as OD_600_ with a spectrophotometer (Genesys 20, Thermo Fisher Scientific). In the minimal MOPS-glucose medium, one OD_600_*ml is equivalent to 0.5 mg cell dry weight (74). To maintain sufficient aeration, 3 mL and 10 mL cultures were grown in 16 and 25 mm diameter glass vessels, respectively, depending on the required volume for each experiment. Growth rate and viability differences between 3 mL and 10 mL culture were not distinguishable from noise.

To achieve reproducible exponential growth prior to starvation, we followed a standard three-stage culturing approach: seed culture, pre-culture and experimental culture. Cells from a single colony on an LB agar were inoculated in LB broth medium and grown for about 4-6 hours to late exponential or early stationary phases (seed-culture). The seed-culture was diluted approximately 10^3^ to 10^5^-fold into MOPS medium supplemented with 10 mM glucose and grown overnight (pre-culture). The pre-culture was used to prepare a main culture according to the procedure described for each experiment.

### Protein degradation extent

Protein degradation extent during starvation was measured by labeling cells during growth with ^14^C-leucine and chasing its release during starvation into the medium containing excess amount of cold leucine as previously described (10). Briefly, cells from pre-culture were washed once and suspended in MOPS medium supplemented with 2.5 mM glucose. The cell suspension was diluted in MOPS-medium supplemented with 2.5 mM glucose and 100 μM L-[^14^C(U)]-leucine (6.7μCi/μmol, CAS No. 61-90-5, Revvity) to OD_600_ = 0.01 and grown to an OD_600_ = 0.4. The cells were harvested at 9,000 rpm (7,607 x g) for 2 minutes (Centrifuge 5424, Eppendorf), washed once with a chase medium (glucose-free MOPS-medium containing 0.81 mM sodium acetate and 1 mM L-[^12^C]-leucine), and suspended in the chase medium to an OD_600_ = 0.4. Acetate was supplemented to make up for the acetate released during growth, which is available in the glucose-runout experiments used for proteomics sampling and viability measurements. The onset of starvation is defined as when the chase medium was added to the harvested cells for washing. At various times after starvation, 100 μl of the culture was withdrawn and precipitated with 200 μl of 20% trichloroacetic acid (TCA) for at least 30 minutes on ice. Half of the mixture was directly transferred to 2 ml scintillation cocktail (Liquiscint, National Diagnostics). The other half was centrifuged at 15,000 rpm (21,130 x g) for 15 minutes (Centrifuge 5424, Eppendorf) and the supernatant was transferred to the scintillation cocktail. The radioactivity was monitored by scintillation. Degradation extent is defined as the radioactivity in the TCA-soluble fraction divided by the total radioactivity of the culture.

### Proteomics

Data for **Figure 1C-E** were obtained from (10). Proteomics data on wild type cells and SPD mutants for **Figure 2C-D** (described in **Table S6**) were collected and analyzed as previously described (10). Briefly, cells were lysed in trifluoroacetic acid and subjected to proteolysis with trypsin. After desalting and washing, peptides were analyzed on a Vanquish Neo (micro-flow configuration) coupled to an Orbitrap Exploris 480 mass spectrometer (both Thermo Fisher Scientific) operated in data-independent acquisition (DIA) mode. Raw files were analyzed with DIA-NN version 2.3 using the UniProt reference fasta file for *E. coli* (K-12, MG1655) UP000000625, and using the same settings as in (10). The main DIA-NN report parquet files were used for all subsequent calculations, and proteins were filtered to a Lib.PG.Q.Value<0.001 with at least two detectable peptides per protein.

Protein abundances were quantified as previously described (10). Briefly, peptide intensities were filtered for Q-values below 10^-3^ and protein intensities were quantified with xTop v2.5 (76) (https://gitlab.com/mm87/xtop) with settings N_c_thres=6 and s2_min=0.03; all other settings were left as default. The std-log errors generated by xTop were used to filter out noisy proteins with std-log errors above 0.3 or at least 2.5x larger than the smallest error across all samples in the batch. Absolute protein mass fractions were computed by calibrating the average protein mass fractions obtained for *E. coli* EQ353 exponentially growing in glucose minimal media to ribosome profiling data obtained for the same exact strain and condition (77); quickly degraded proteins or proteins for which we suspect biological differences across samples were not rescaled as done previously (10). Protein levels during the runout experiments were expressed as mass fraction of total protein mass and are provided in **Table S7**.

### Bioinformatics

For analysis of operon structure, the sequence of *E. coli* SpdB/YeaH (UniProt ID P26235) was used as a query for the *fast.genomics* server (22), and 200 “random hits” were plotted from a total of 1,656 hits identified by mmseqs2 (78). A 12-kb region around each hit was plotted, with gene clusters conserved in at least 30% of operons colored. For the SpdB phylogenetic tree, a sequence alignment of YeaH homologs was performed in MUSCLE version 3 (79) using fast settings (-maxiters 2 -maxmb 1000), and the tree was generated with FastTree 2 (80) using default settings, using aligned regions of identified SpdB homologs. All proteins identified in this analysis and used for sequence alignments of SpdA, SpdB, and SpdC homologs are listed in **Table S1**.

### Protein structure predictions

For AlphaFold 2 protein interaction screens, AlphaPulldown (81) version 1.0.4 was used to perform a scan of SpdA, SpdB and SpdC against a list of 42 protein degradation-associated proteins and σ^S^ regulon proteins, including SpdA, SpdB and SpdC (**Table S2**). Briefly, protein features were first calculated using ‘create_individual_features.py’ with mmseqs2 (78) (‘-- use_mmseqs2=True’) and storing the MSA files. Then, protein pairs were analyzed with ‘run_multimer_jobs.py’ in pulldown mode (‘--mode=pulldown’) by providing the directory for the monomer features. Other settings include ‘--use_hhsearch=True’, ‘--num_cycle=2’, ‘-- num_predictions_per_model=1’, ‘model_to_relax=None’.

For structure predictions using AlphaFold 3 (31), the AlphaFold Server (https://alphafoldserver.com) was used with standard settings. Predictions involving SpdA (or other MoxR-type ATPases) included one ATP and one Mg^2+^ ion per SpdA protomer; predictions involving SpdB (or other vWA domain proteins) included one Mg^2+^ per SpdB protomer; and predictions involving SpdC included one Zn^2+^ ion per SpdC protomer. Structure comparisons were performed with the DALI server (32). For visualization of predicted aligned error (PAE) plots, AlphaFold model (.cif) and confidence (.json) files were uploaded to the PAE Viewer web site (https://pae-viewer.uni-goettingen.de) (82). All protein sequences used for structure predictions are listed in **Table S3**.

### Protein expression and purification

For protein purification, genes were individually cloned into *E. coli* expression vectors encoding either no tag (UC Berkeley Macrolab vector 2A-T, Addgene ID 29665), an N-terminal TEV protease-cleavable His_6_-tag (UC Berkeley Macrolab vector 2B-T, Addgene ID 29666), or an N- terminal His_6_-maltose binding protein (MBP)-TEV-tag (UC Berkeley Macrolab vector 2C-T, Addgene ID 29706). For SpdB and SpdC co-expression, a C-terminal His_6_-SUMO-tagged SpdB and untagged SpdC were generated by PCR and cloned into UC Berkeley Macrolab vector 2A-T. Targeted mutations were generated by PCR-based mutagenesis.

Vectors were transformed into *E. coli* Rosetta 2 pLysS (EMD Millipore), and 1L cultures were grown at 37°C in 2XYT media to an OD_600_ of 0.6 before induction with 0.25 mM IPTG at 20°C for 16-18 hours. Cells were harvested by centrifugation, resuspended in buffer A (25 mM Tris-HCl pH 8.5, 300 mM NaCl, 5 mM MgCl_2_, 10% glycerol and 5 mM β-mercaptoethanol) containing 5 mM imidazole, then lysed by sonication (Branson Sonifier). Lysates were clarified by centrifugation, then supernatants were passed over a Ni-NTA Superflow column (Qiagen) pre-equilibrated in buffer A plus 5 mM imidazole. The column was washed in buffer A containing 20 mM imidazole, then bound protein was eluted in buffer A containing 400 mM imidazole. Eluates were concentrated by ultrafiltration (Amicon Ultra; EMD Millipore). Proteins were further passed over a Superdex 200 Increase size exclusion column (Cytiva) in size exclusion buffer (25 mM Tris-HCl pH 8.5, 300 mM NaCl, 5 mM MgCl_2_, 10% glycerol, and 1 mM DTT) or a Superose 6 Increase size exclusion column (Cytiva) in the same buffer. Peak fractions were concentrated by ultrafiltration and stored at -80°C for biochemical assays.

### SEC-MALS

For size exclusion chromatography coupled to multi-angle light scattering (SEC-MALS), 100 µl of purified proteins at a concentration of 2 mg/ml were injected onto a size exclusion column (Superdex 200 Increase 10/300 GL, Cytiva) in size exclusion buffer, then light scattering and differential refractive index (dRI) profiles were collected using miniDAWN TREOS and Optilab T-rEX detectors (Wyatt Technology). SEC-MALS data were analyzed using ASTRA software version 8 and visualized with Prism version 10 (GraphPad Software).

### Yeast two-hybrid assays

Genes of SpdA, SpdB and SpdC were individually cloned into vectors pBridge (BD fusion) and pGADT7 (AD fusion;Clontech). Plasmids were transformed into *S. cerevisiae* strains AH109 (for AD fusions) and Y187 (for BD fusions) and selected on single-dropout media lacking leucine or tryptophan. Selected AD/BD pairs were mated and selected on double-dropout media (-LT; lacking leucine and tryptophan). For interaction tests, cells were resuspended and equal numbers were plated onto non-selective (-LT), selective (-LTH; lacking leucine, tryptophan, and histidine), and stringently selective (-LTHA, lacking leucine, tryptophan, histidine, and adenine) plates, grown overnight at 30°C, then imaged.

## ACKNOWLEDGEMENTS

This work was supported by the NIH (R35GM152133 to T.H., R35GM144121 to K.D.C.), the NSF (MCB2545232 to T.H.), and by the Horizon 2020 program of the European Union (EPIC-XS, Project No. 823839 to C.L.). The Exploris 480 mass spectrometer used in this study was funded in part by the German Research Foundation (INST 95/1435-1 FUGG). Additionally, G.M. acknowledges the Department of Defense (DOD) National Defense Science and Engineering Graduate (NDSEG) Fellowship Program, L.R.C. acknowledges NIH T32GM139795 and F31AI194603, and C.L. and T.H. acknowledge support by the Bavaria California Technology Center (BaCaTeC, Project No. 2023-1).

---

**Figure S1.**
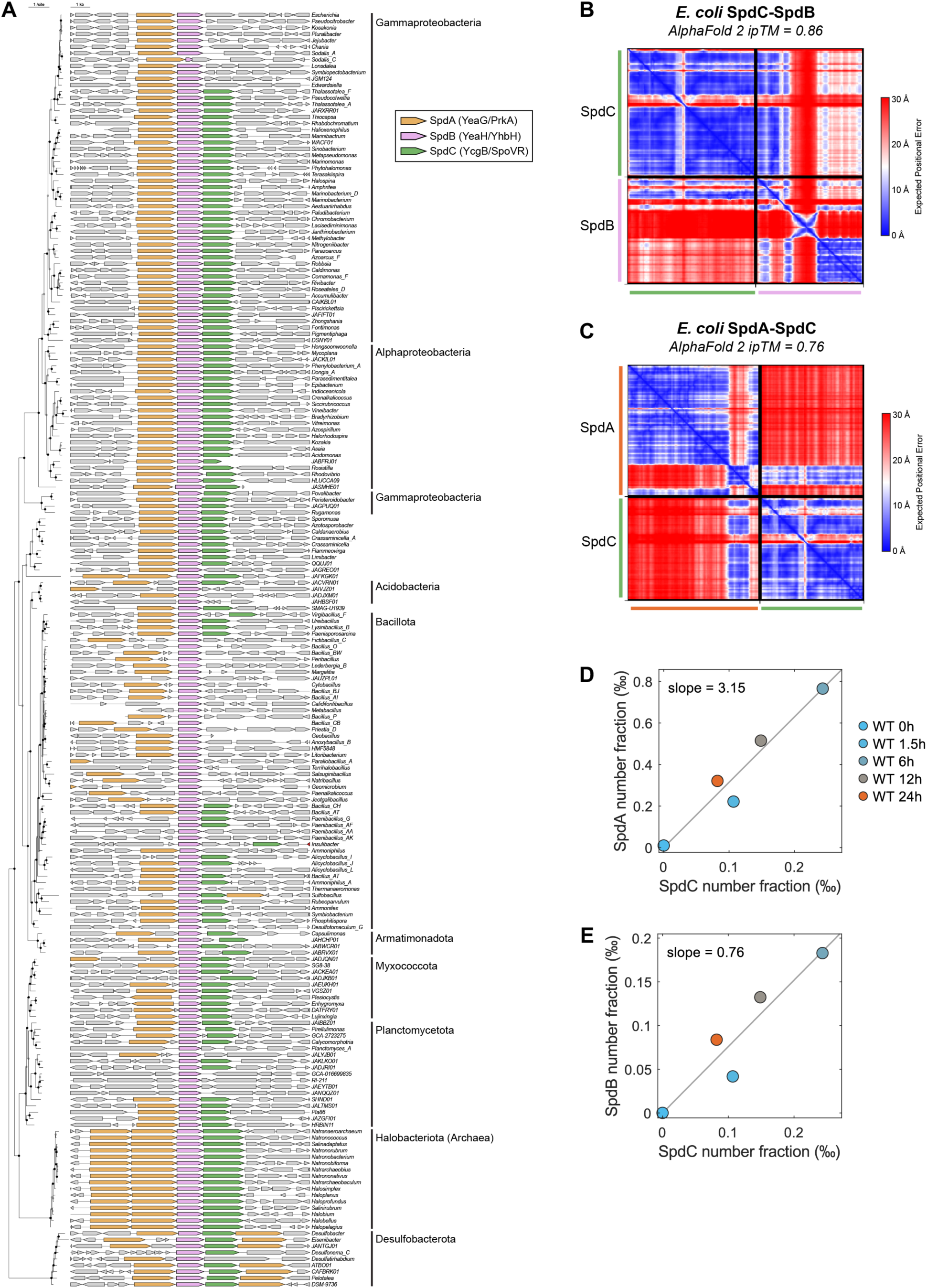
Identification of a conserved bacterial stress-response pathway. (A) Gene neighborhoods of selected *spdB* homologs in bacteria and archaea, as identified by the *fast.genomics* server (22). *spdA* homologs are shown in orange, *spdB* pink, and *spdC* green. See **Table S1** for accession numbers. Phylogenetic tree calculated from an alignment of each YeaH protein sequence. (B) AlphaFold 2 predicted aligned error (PAE) plot for *E. coli* SpdC and SpdB. See **Table S2** for full AlphaFold 2 screening results. (C) AlphaFold 2 predicted aligned error (PAE) plot for *E. coli* SpdA and SpdC. (D) Comparison of the copy number fraction (mass fraction corrected for each protein’s molecular weight) of SpdA versus SpdC, from quantitative mass spectrometry data in glucose-starved wild type (WT) cells (10). (E) Comparison of the copy number fraction of SpdB versus SpdC, from quantitative mass spectrometry data in glucose-starved wild type (WT) cells (10). Using the slopes calculated in panels (D) and (E), the inferred molar ratio is 4.14 SpdA:1 SpdB:1.32 SpdC (reported as 4:1:1 in the main text).

**Figure S2.**
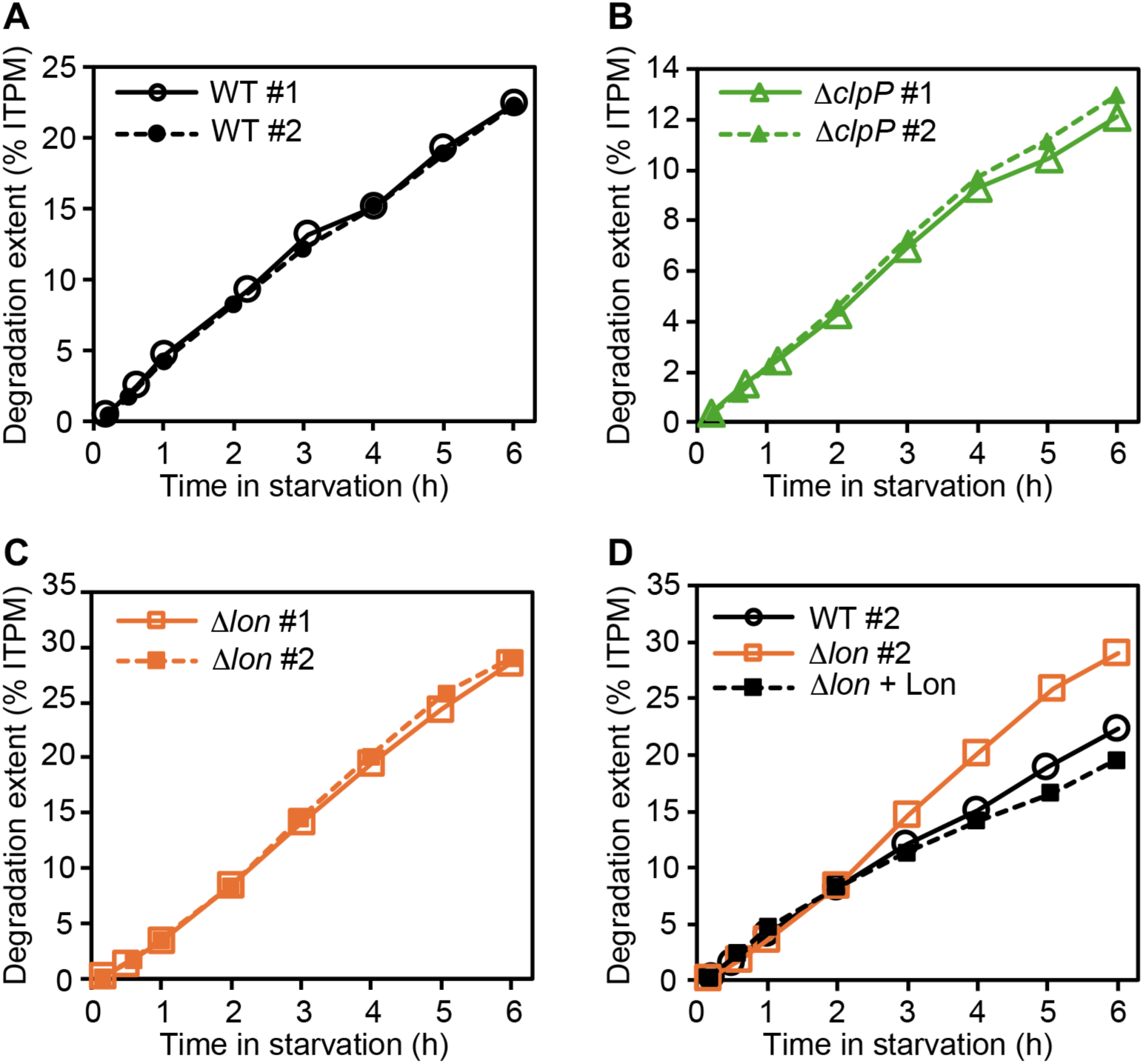
Repeatability of degradation extent measurements. (A-C) Degradation extent measurements are highly repeatable. Shown in these panels are repeats done for various strains, with some repeats done months apart using different batches of ^14^C-leucine. The repeats are not shown in the main result figures for their ease of readability. (D) Complementation of Lon in a strain harboring *lon* expression from a plasmid in Δ*lon* background (filled squares).

**Figure S3.**
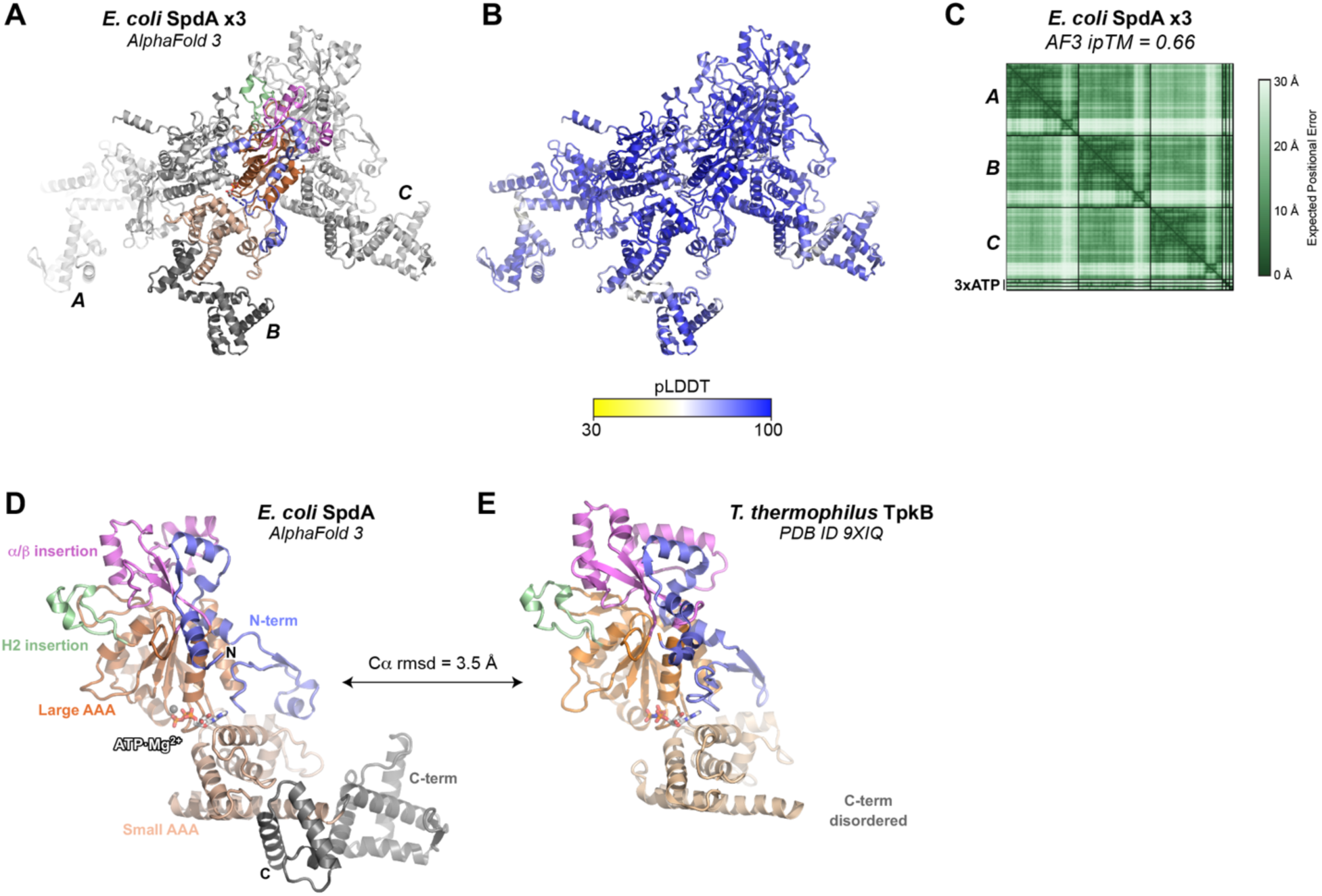
AlphaFold 3 predictions of *E. coli* SpdA. (A) AlphaFold 3 predicted structure of *E. coli* SpdA (three copies, protomer B colored as in Figure 3B while protomers A and C are colored gray), plus three ATP molecules and three Mg^2+^ ions. (B) View as in panel (A), colored by local confidence (predicted Local Distance Difference Test; pLDDT). (C) AlphaFold 3 predicted aligned error (PAE) plot for the *E. coli* SpdA structure prediction. (D) View of one *E. coli* SpdA protomer, colored as in Figure 3B. Bound ATP is shown in sticks, and Mg^2+^ ion is shown as a gray sphere. (E) View of *T. thermophilus* TpkB (PDB ID 9XIQ) (15) aligned to SpdA in panel (D), and with domains colored as in SpdA. Bound AMPPNP is shown in sticks.

**Figure S4.**
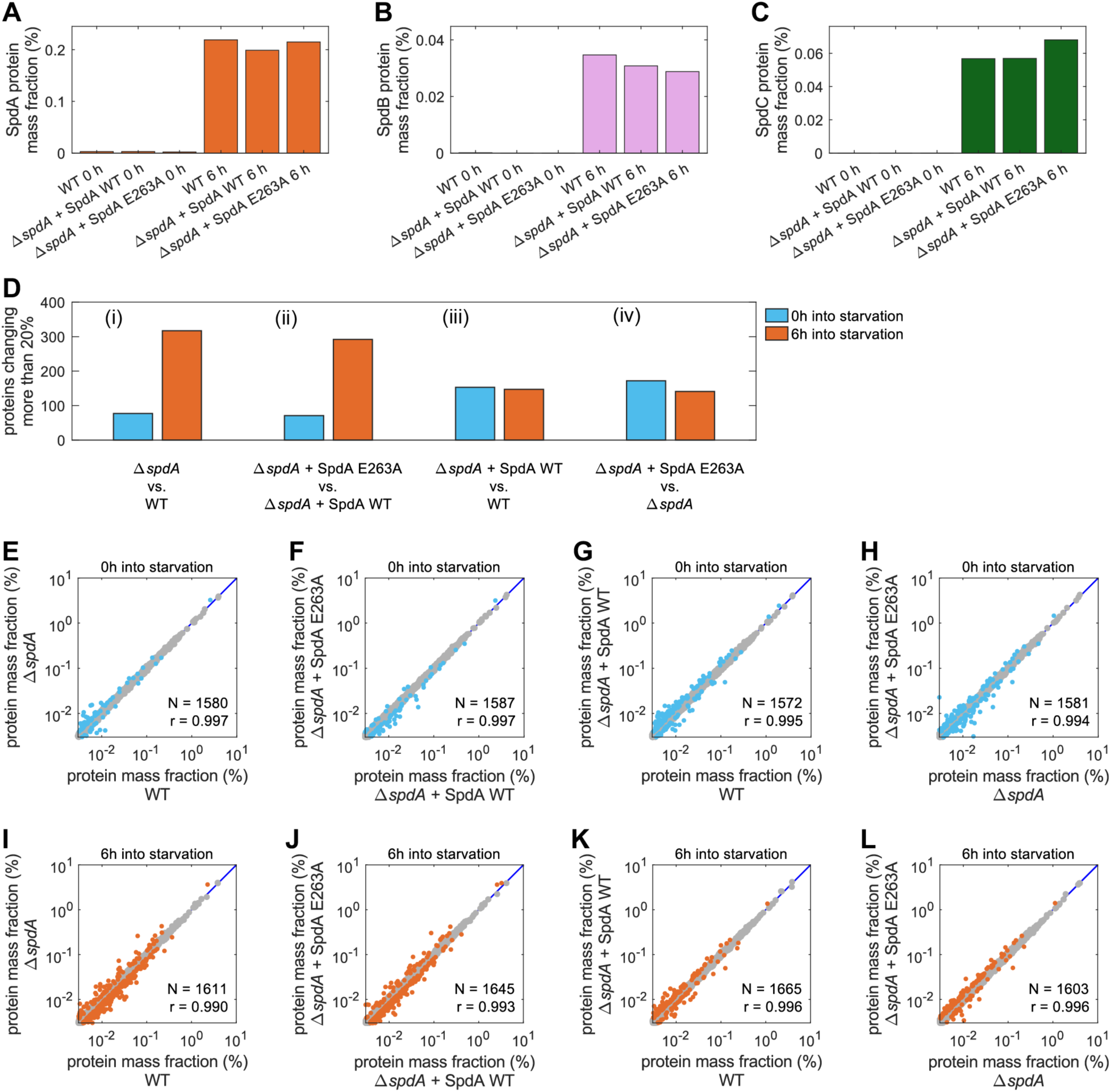
SpdA E263A mutant and its wild type complement. (A) SpdA protein mass fraction during exponential growth (0 h) and at 6h after glucose depletion, for wild type (WT) and Δ*spdA* complemented with plasmids encoding wild type SpdA or the E263A point mutant. (B) SpdB protein mass fraction during exponential growth (0 h) and at 6h after glucose depletion, in the same strains as panel (A). (C) SpdC protein mass fraction during exponential growth (0 h) and at 6h after glucose depletion, in the same strains as panel (A). (D) Summary of pairwise proteome comparisons (see panels E-L), showing the number of proteins where the absolute abundance changes by more than 20% between the two indicated strains. Blue bars indicate comparisons for the proteomes during exponential growth (0 h) and orange bars indicate comparisons 6 h after glucose depletion. Four pairwise comparisons are made: (i) Δ*spdA* cells versus wild type (WT) cells; (ii) Δ*spdA* + SpdA E263A mutant and Δ*spdA* + WT SpdA; (iii) Δ*spdA* + WT SpdA and WT; (iv) Δ*spdA* + SpdA E263A mutant and Δ*spdA*. Several observations can be made from these comparisons: (1) The similarity of comparisons (i) and (ii) shows that expression of SpdA E263A does not complement Δ*spdA* as measured by the change in proteome after starvation. Comparison (iv) further shows that the changes due to *spdA* deletion and the E263A mutations are similar. (2) Comparisons (iii) and (iv) show that the plasmid harboring either WT or E263A mutant of *spdA* does not induce the large changes between 0 and 6h seen in comparison (ii). (3) Comparing the blue bars (exponential growth) of the pairs (i) and (iii) gives a glimpse of the subtle effect of the plasmid harboring WT *spdA*. The only difference in this comparison is the plasmid harboring WT *spdA* which is driven by the native spdA promoter. During exponential growth, SpdA is not expected to be expressed, hence little difference is seen between Δ*spdA* and WT cells. However, some moderately increased differences are seen between WT cells and Δ*spdA* with WT SpdA. This suggests that the plasmid harboring SpdA has some subtle effect already on exponentially growing cells, e.g., from leaky expression of SpdA. (E-L) Scatter plots showing protein abundance comparisons summarized in panel (D).

**Figure S5.**
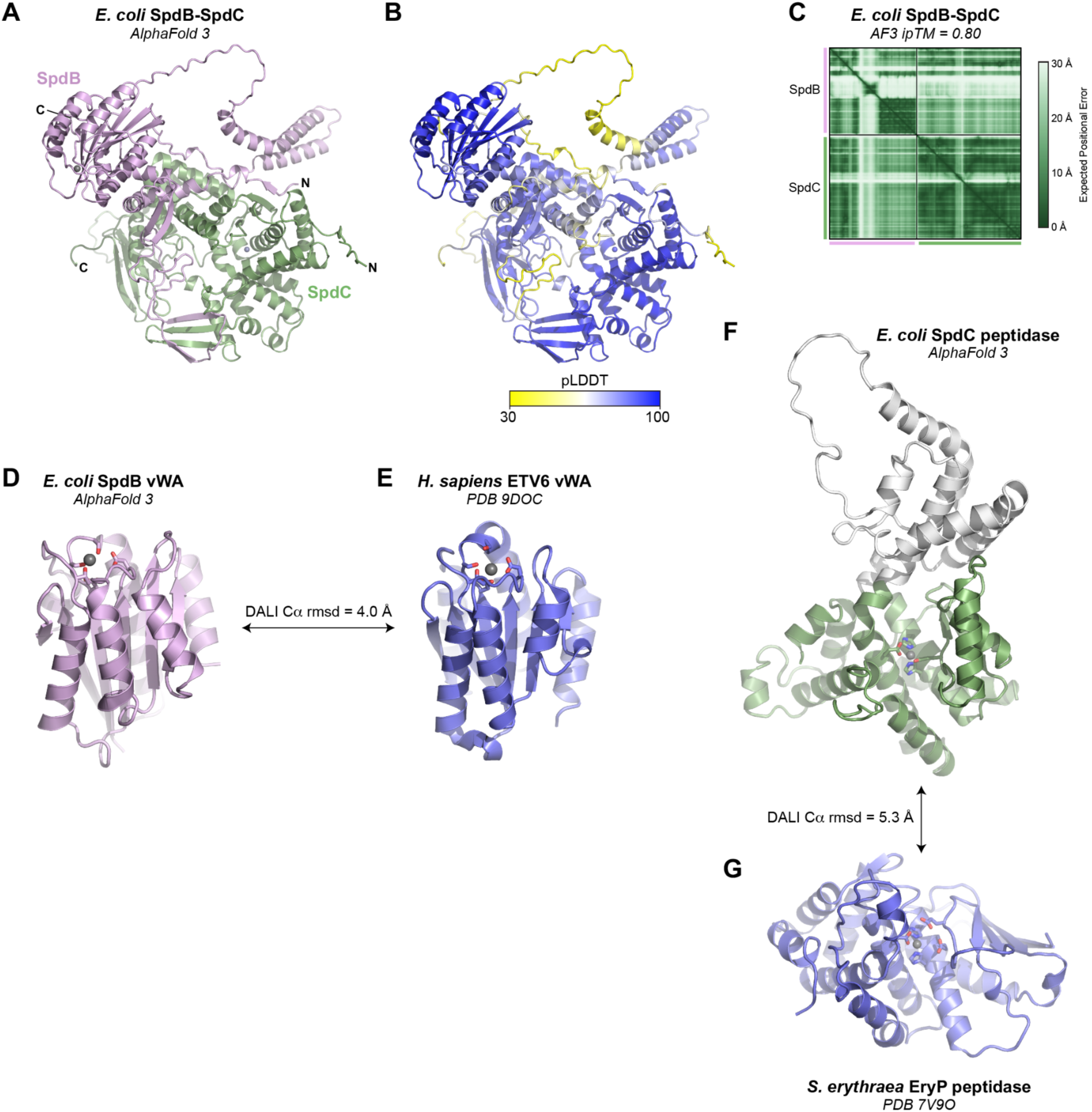
AlphaFold 3 structure prediction of the *E. coli* SpdB-SpdC complex. (A) Overall structure of *E. coli* SpdB-SpdC predicted by AlphaFold 3, with SpdB shown in pink and SpdC in green. (B) Overall structure of *E. coli* SpdB-SpdC predicted by AlphaFold 3, colored by local prediction confidence (pLDDT). (C) AlphaFold 3 predicted aligned error (PAE) plot for *E. coli* SpdB-SpdC. (D) View of the predicted vWA domain of *E. coli* SpdB bound to Mg^2+^. (E) View of *H. sapiens* capillary morphogenesis gene II vWA domain (PDB 9DOC; unpublished) bound to Mg^2+^. A DALI superposition of this domain with *E. coli* SpdB gives an overall Cα r.m.s.d. of 4.0 Å. (F) View of the predicted M1 zinc metallopeptidase domain of *E. coli* SpdC bound to Zn^2+^, with insertion domain (residues 135-273) shown in white. (G) View of *S. erythraea* EryP M1 zinc metallopeptidase domain bound to Zn^2+^ (PDB ID 7V9O) (86). A DALI superposition of this domain with *E. coli* SpdC gives an overall Cα r.m.s.d. of 5.3 Å (see **Table S4**).

**Figure S6.**
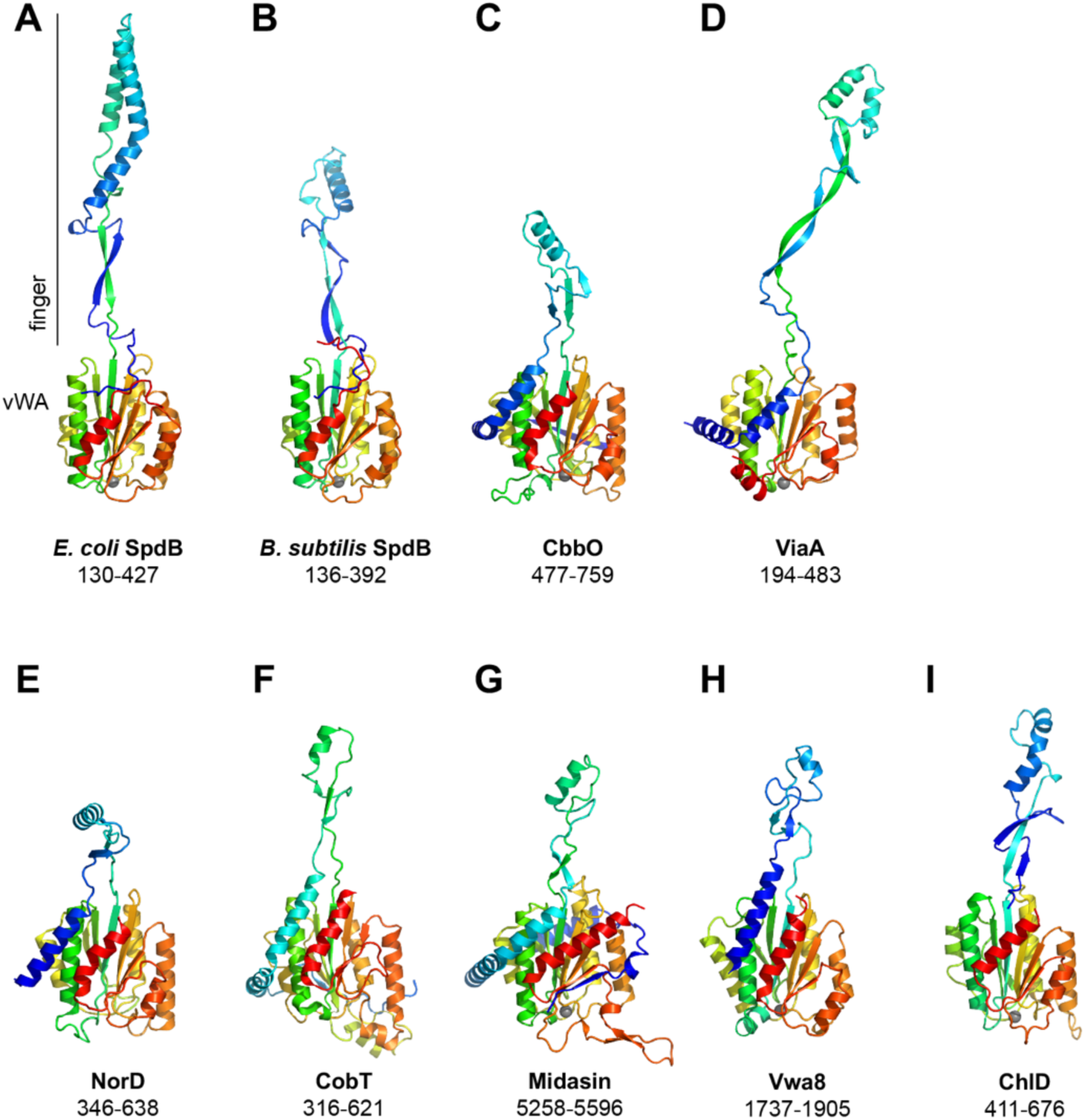
Shared architecture of vWA proteins in chaperone systems. (A) Finger-vWA region of *E. coli* SpdB from an AlphaFold prediction of an SpdA-SpdB-SpdC complex (**Figure S7**), colored as a rainbow from residue 130 (blue) to 427 (red). Bound Mg^2+^ ion is shown as a gray sphere. The finger and vWA domains are labeled. (B) Finger-vWA region of *B. subtilis* SpdB from an AlphaFold prediction of an SpdA-SpdB-SpdC complex (**Figure S8**), colored as a rainbow from residue 136 (blue) to 392 (red). (C) Finger-vWA region of *Acidithiobacillus ferrooxidans* CbbO2 from an AlphaFold prediction of a CbbQ2-CbbO2 complex (**Figure S9**), colored as a rainbow from residue 477 (blue) to 759 (red). (D) Finger-vWA region of *E. coli* ViaA from an AlphaFold prediction of a RavA-ViaA complex (**Figure S10**), colored as a rainbow from residue 194 (blue) to 483 (red). (E) Finger-vWA region of *Paracoccus denitrificans* NorD from a cryoEM structure of NorQ-NorD complex (PDB 9QH8) (33), colored as a rainbow from residue 346 (blue) to 638 (red). (F) Finger-vWA region of *Brucella melitensis* CobT from a cryoEM structure of CobS-CobT (PDB 9UKG) (34), colored as a rainbow from residue 316 (blue) to 621 (red). (G) Finger-vWA region of *H. sapiens* Midasin from an AlphaFold prediction of the Midasin AAA+ and finger-vWA domains (**Figure S14**), colored as a rainbow from residue 5258 (blue) to 5596 (red). (H) Finger-vWA region of *H. sapiens* Vwa8 from an AlphaFold prediction of full-length Vwa8 (**Figure S15**), colored as a rainbow from residue 1737 (blue) to 1905 (red). (I) Finger-vWA region of *Synechocystis* ChlD from AlphaFold prediction of a ChlI-ChlD-ChlH complex (**Figure S11**), colored as a rainbow from residue 411 (blue) to 676 (red).

**Figure S7.**
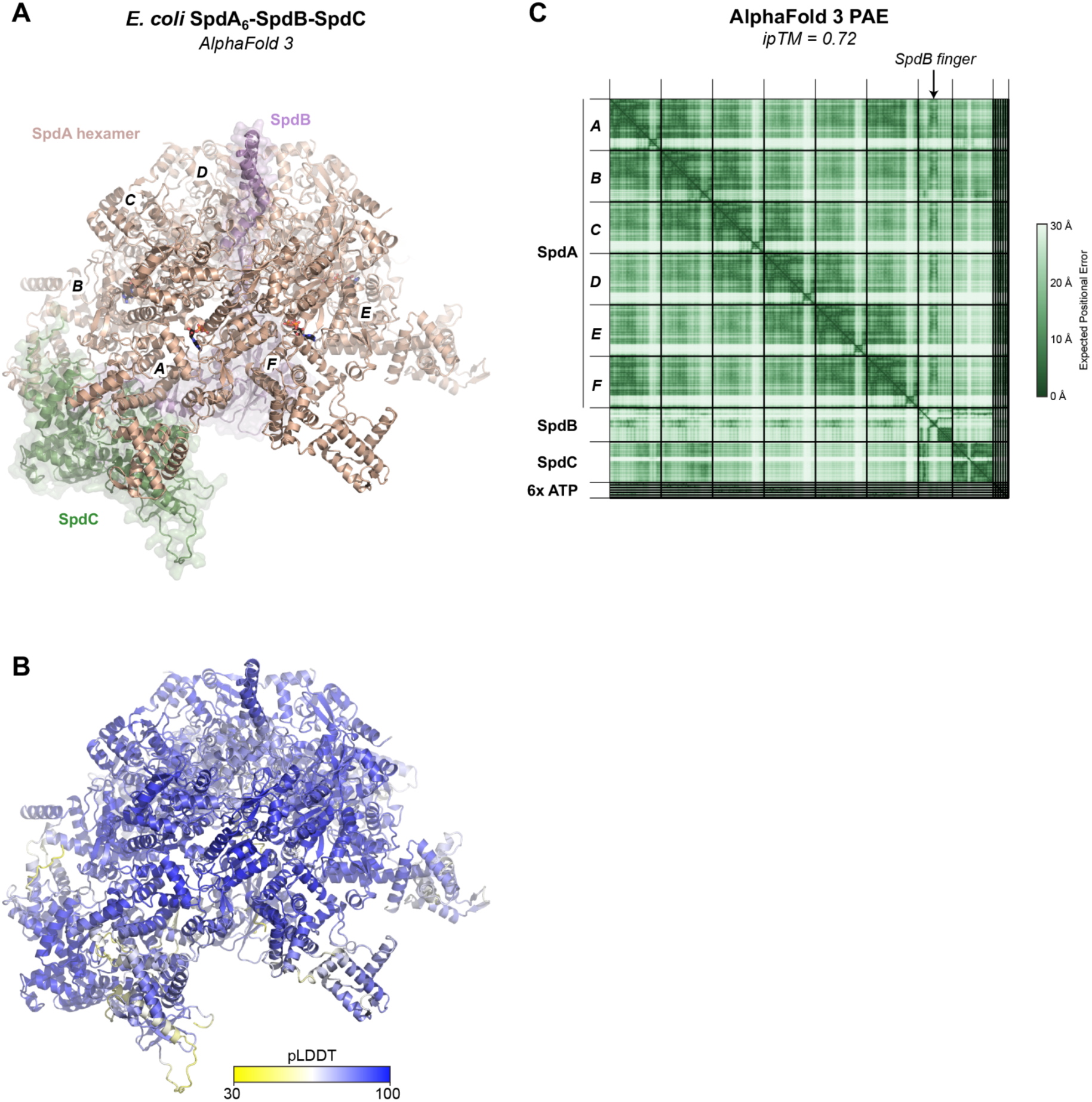
AlphaFold prediction of *E. coli* SpdA-SpdB-SpdC. (A) AlphaFold 3 predicted structure of *E. coli* SpdA (six copies, colored light brown with protomers A-F labeled), SpdB (pink), and SpdC (green). Also included in the prediction were six ATP molecules, seven Mg^2+^ ions, and one Zn^2+^ ion. (B) View as in panel (A), colored by local confidence (pLDDT). (C) AlphaFold 3 predicted aligned error (PAE) plot for the *E. coli* SpdA-SpdB-SpdC structure prediction. The location of the SpdB finger domain is indicated with an arrow at top.

**Figure S8.**
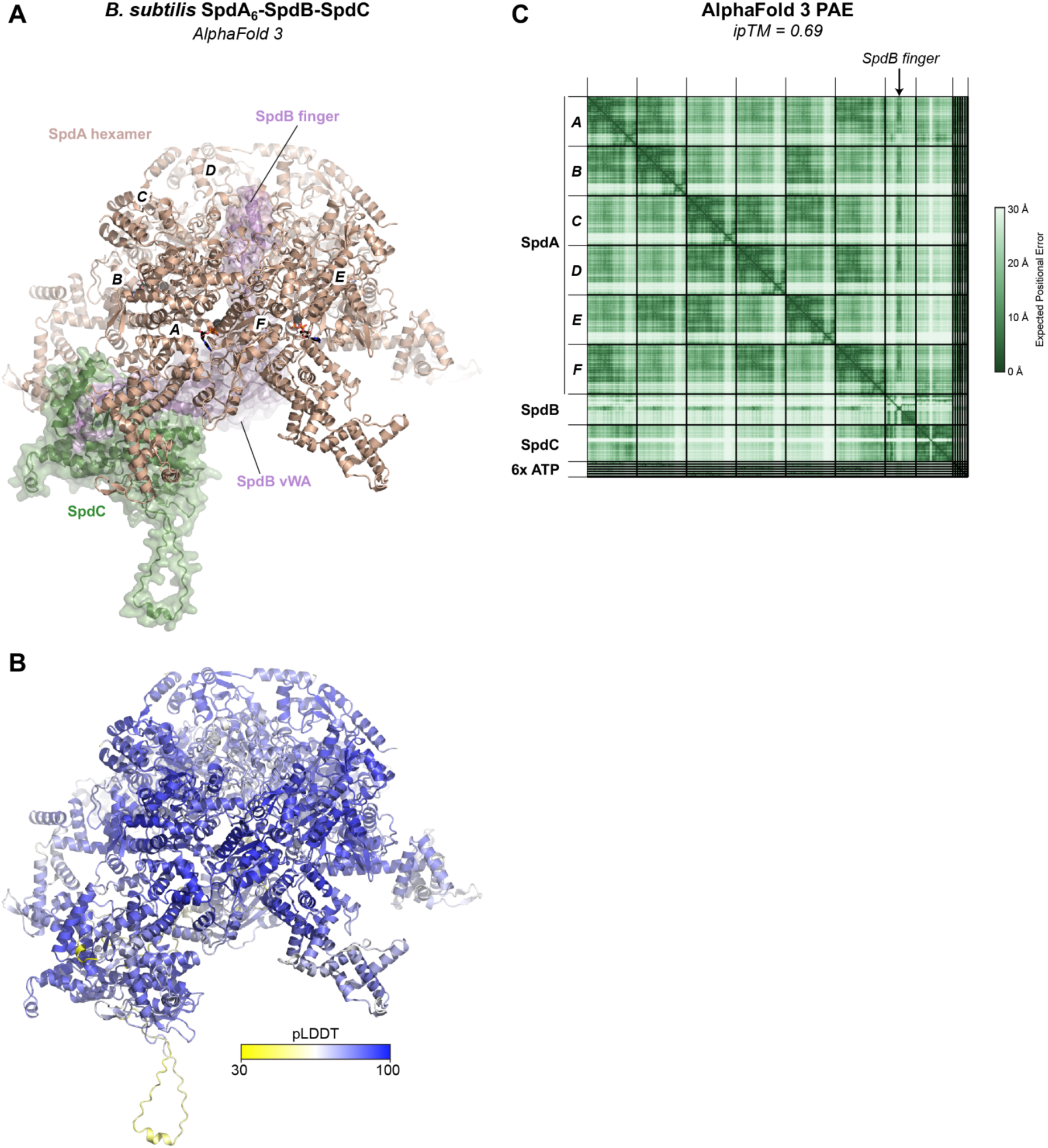
AlphaFold prediction of *B. subtilis* SpdA-SpdB-SpdC. (A) AlphaFold 3 predicted structure of *B. subtilis* SpdA (six copies, colored light brown with protomers A-F labeled), SpdB (pink), and SpdC (green). Also included in the prediction were six ATP molecules, seven Mg^2+^ ions, and one Zn^2+^ ion. (B) View as in panel (A), colored by local confidence (pLDDT). (C) AlphaFold 3 predicted aligned error (PAE) plot for the *B. subtilis* SpdA-SpdB-SpdC structure prediction. The location of the SpdB finger domain is indicated with an arrow at top.

**Figure S9.**
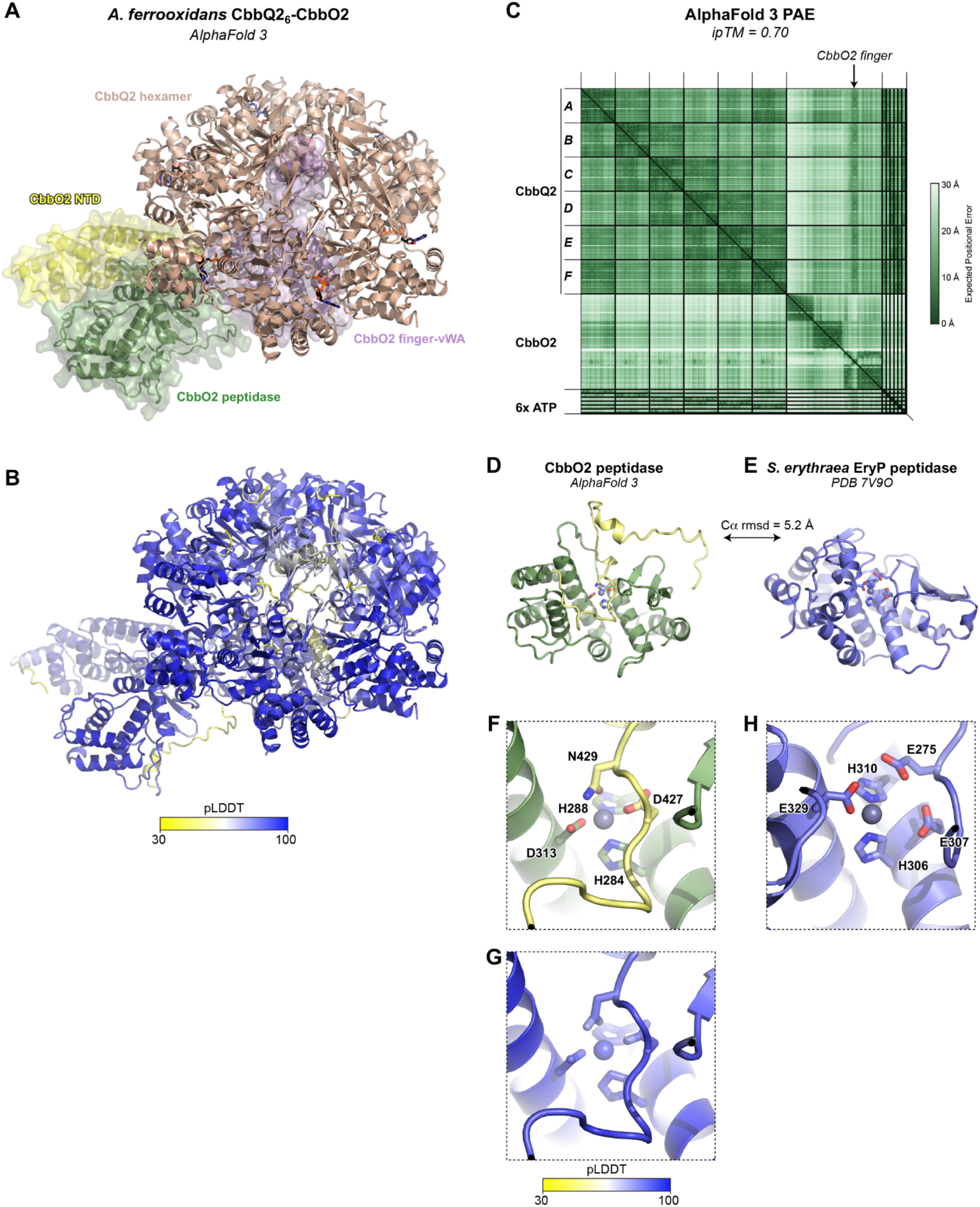
AlphaFold prediction of *A. ferrooxidans* CbbQ2-CbbO2. (A) AlphaFold 3 predicted structure of *A. ferrooxidans* CbbQ2 (six copies, colored light brown) and CbbO2 (N-terminal domain yellow, middle peptidase-like domain green, and finger-vWA domain pink). Also included in the prediction were six ATP molecules, seven Mg^2+^ ions, and one Zn^2+^ ion. (B) View as in panel (A), colored by local confidence (pLDDT). (C) AlphaFold 3 predicted aligned error (PAE) plot for the *A. ferrooxidans* CbbQ2-CbbO2 structure prediction. The location of the CbbO2 finger domain is indicated with an arrow at top. (D) View of the predicted M1 zinc metallopeptidase domain of *A. ferrooxidans* CbbO2, with modeled Zn^2+^ ion shown as a gray sphere. (E) View of *S. erythraea* EryP M1 zinc metallopeptidase domain bound to Zn^2+^ (PDB ID 7V9O) (86). A superposition of this domain with *A. ferrooxidans* CbbO2 gives an overall Cα r.m.s.d. of 5.2 Å. (F) Closeup of the predicted Zn^2+^ binding site of *A. ferrooxidans* CbbO2. Two residues in a confidently-modeled segment C-terminal to the peptidase domain proper (yellow) are predicted to interact with the bound Zn^2+^ ion, suggesting that the peptidase activity may be autoregulated or that the predicted peptidase active site is inactive. (G) View as in panel (F), colored by local confidence (pLDDT). (H) Closeup of the Zn^2+^ binding site of *S. erythraea* EryP M1 zinc metallopeptidase domain (PDB ID 7V9O) (86).

**Figure S10.**
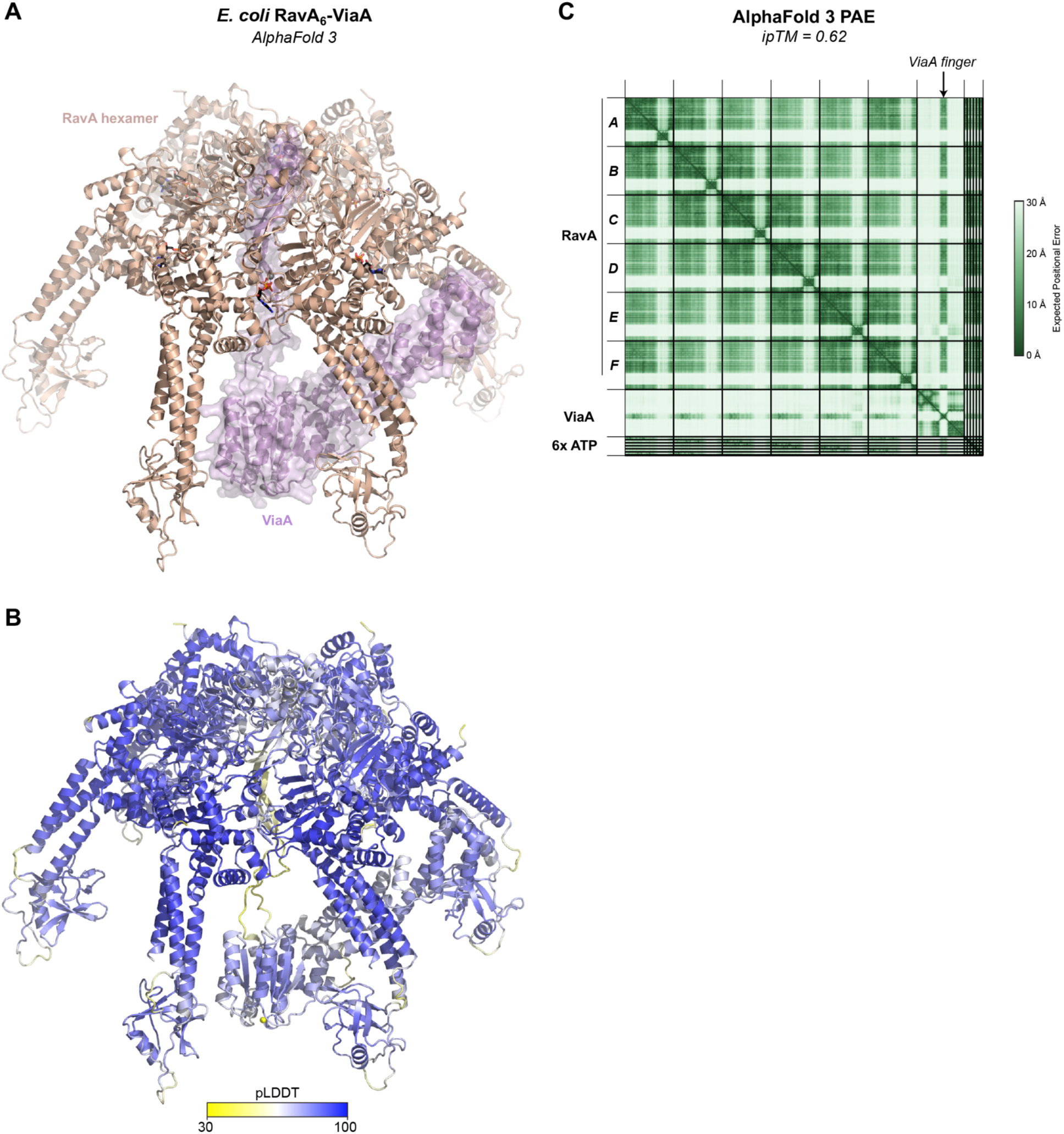
AlphaFold prediction of *E. coli* RavA-ViaA. (A) AlphaFold 3 predicted structure of *E. coli* RavA (six copies, colored light brown) and ViaA (pink). Also included in the prediction were six ATP molecules and seven Mg^2+^ ions. (B) View as in panel (A), colored by local confidence (pLDDT). (C) AlphaFold 3 predicted aligned error (PAE) plot for the *E. coli* RavA-ViaA structure prediction. The location of the ViaA finger domain is indicated with an arrow at top.

**Figure S11.**
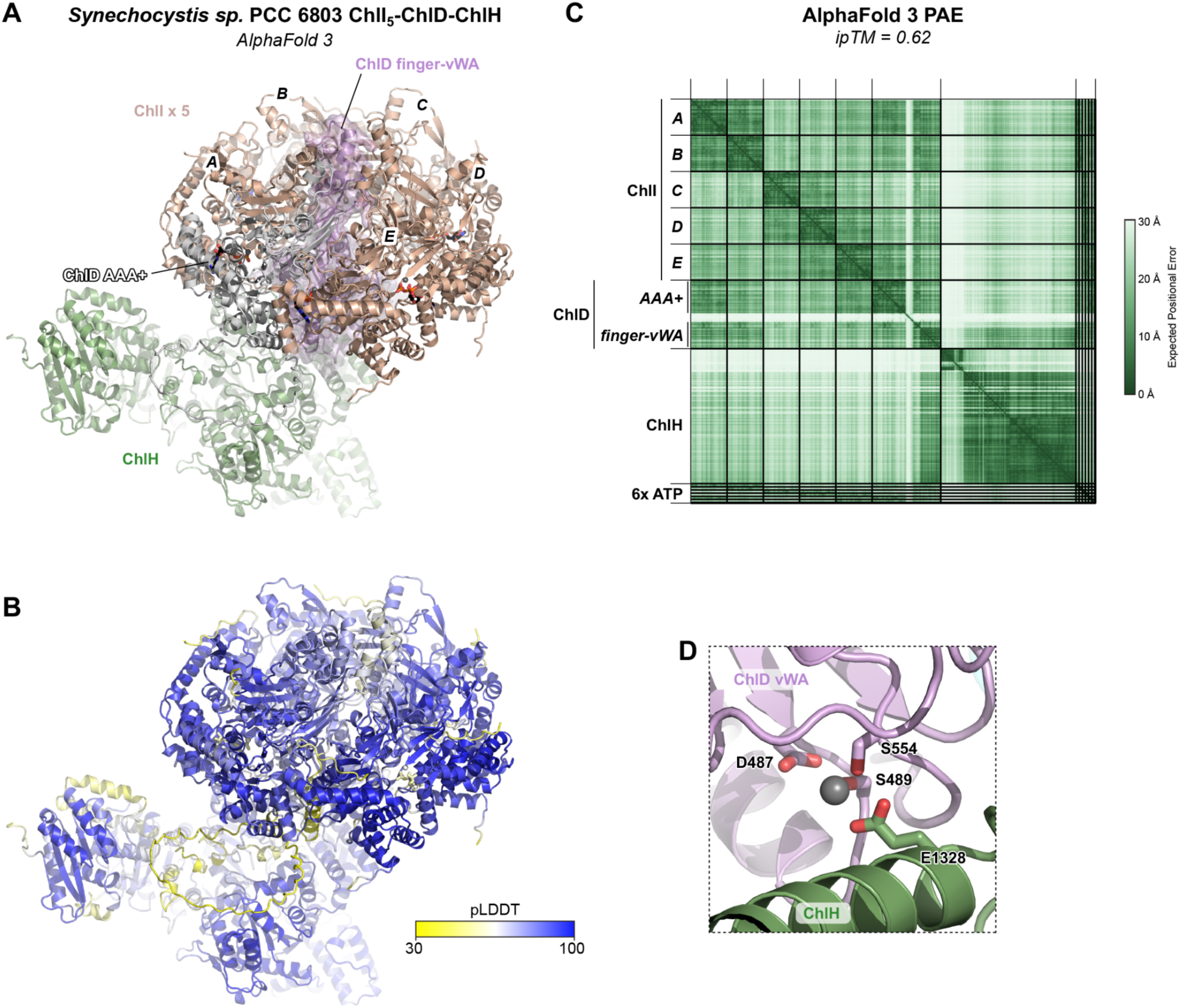
AlphaFold prediction of *Synechocystis* ChlI-ChlD-ChlH. (A) AlphaFold 3 predicted structure of *Synechocystis* sp. PCC6803 ChlI (UniProt ID P51634; five copies, colored light brown with protomers A-E labeled), ChlD (UniProt ID P72772; AAA+ domain white, finger-vWA domain pink), and ChlH (UniProt ID P73020; green). Also included in the prediction were six ATP molecules and seven Mg^2+^ ions. (B) View as in panel (A), colored by local confidence (pLDDT). (C) AlphaFold 3 predicted aligned error (PAE) plot for the *Synechocystis* sp. PCC6803 ChlI-ChlD-ChlH structure prediction. (D) Closeup of the predicted interaction between the ChlD vWA domain (pink with bound Mg^2+^ ion in gray) and ChlH (green).

**Figure S12.**
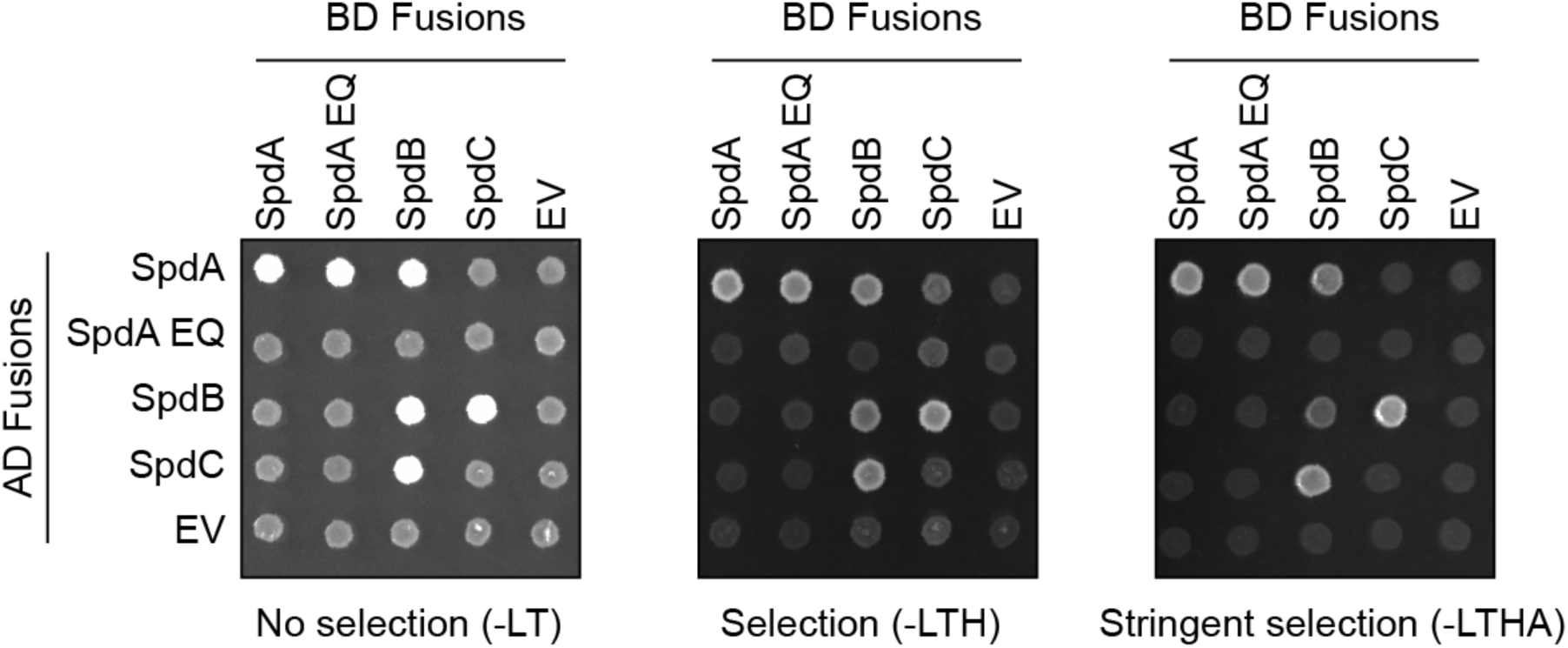
SpdA-SpdB-SpdC yeast two hybrid assays. Yeast two-hybrid results for *E. coli* SpdA (WT or E263Q mutant), SpdB, and SpdC. AD: activation domain; BD: DNA-binding domain; EV: empty vector. Shown is growth on non-selective plates (-LT), selection plates (-LTH), and stringent-selection plates (-LTHA; missing leucine, tryptophan, histidine, and adenine).

**Figure S13.**
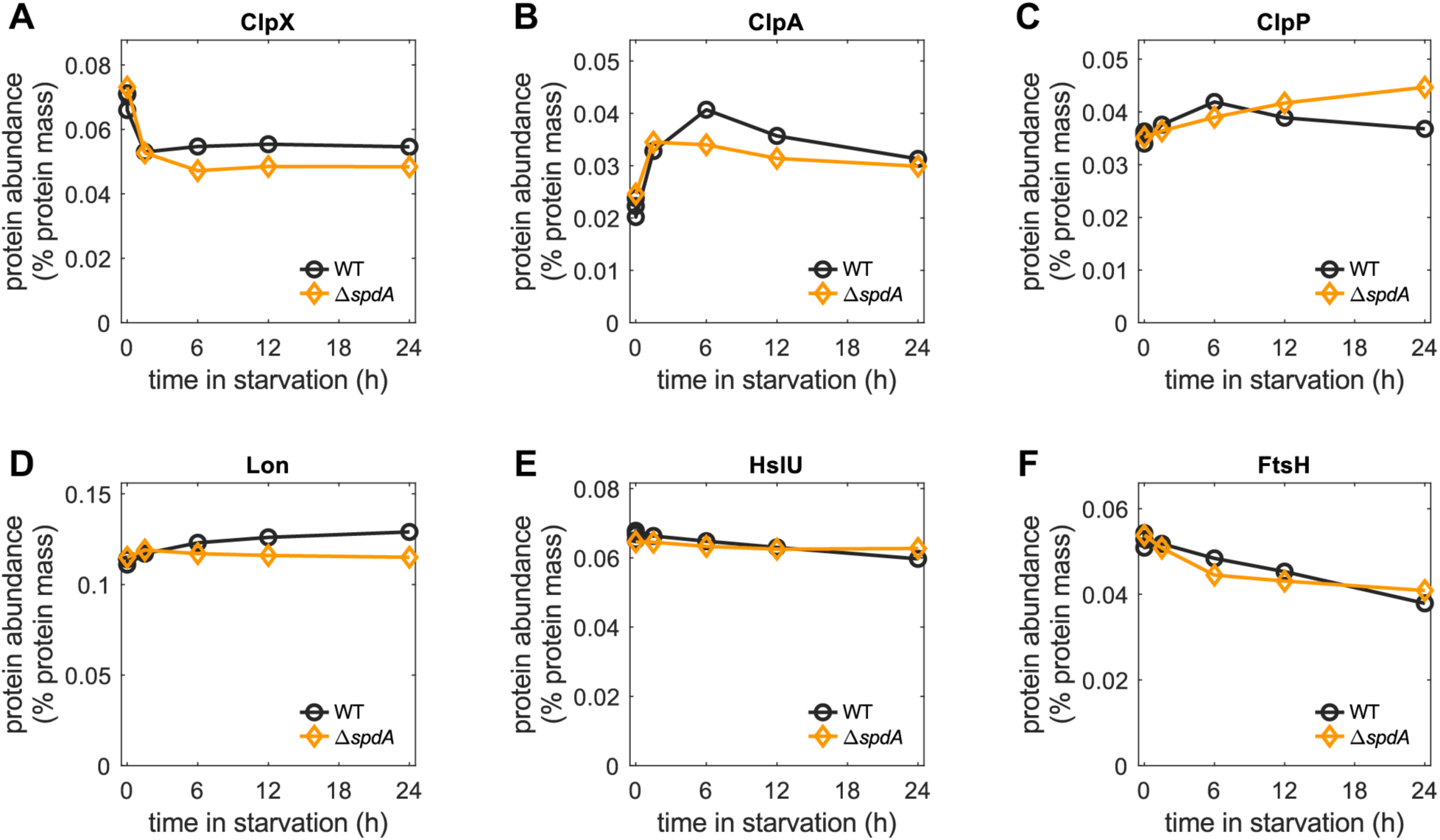
Abundance of cellular proteases in carbon starvation. (A-F) Proteomic data obtained from ref. (10) show that the protein components of the major constitutive proteases, ClpXP, ClpAP, Lon, HslUV, FtsH, remained similar 24 hours into glucose starvation regardless of the presence of the SPD complex.

**Figure S14.**
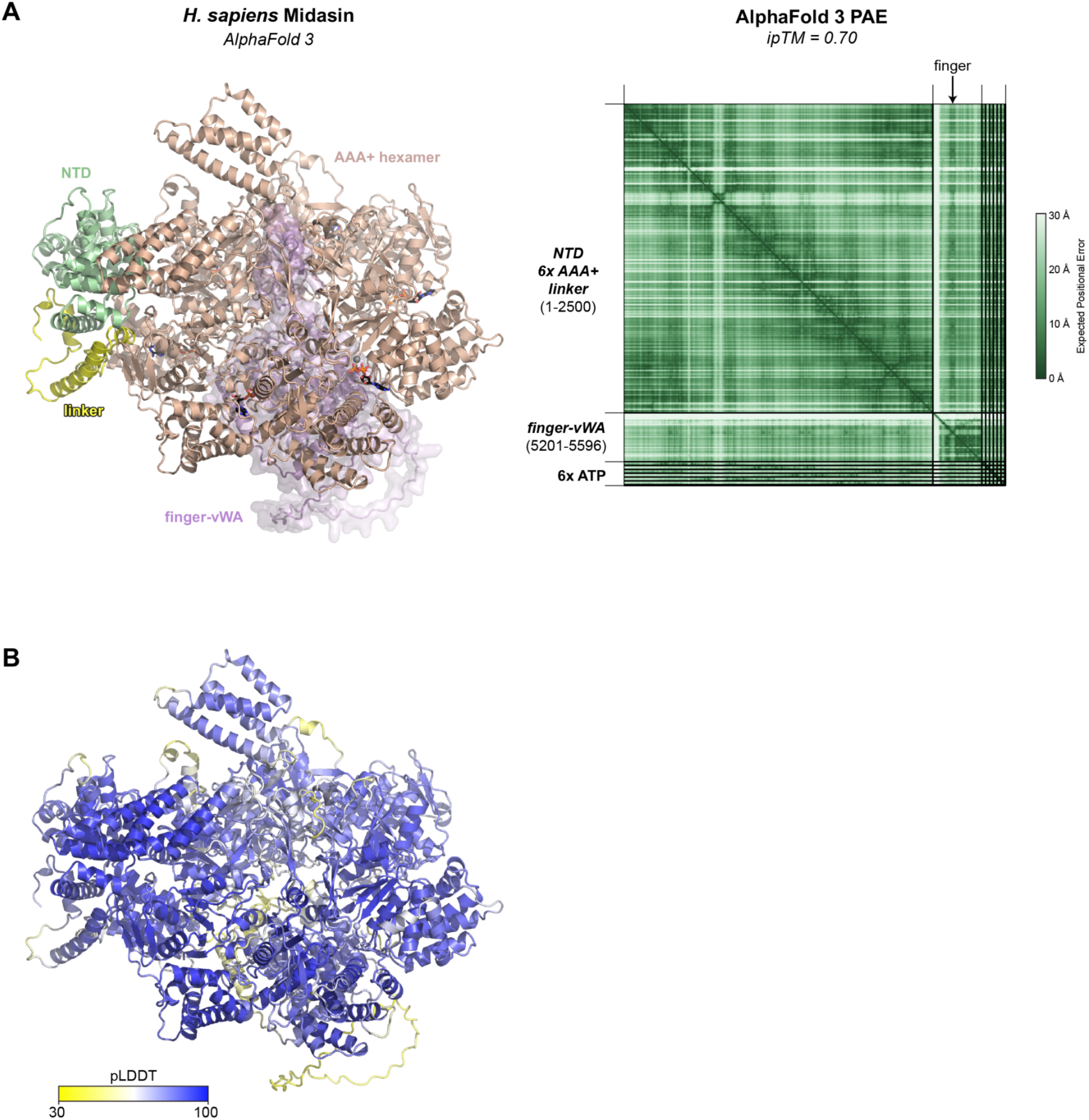
AlphaFold prediction of *H. sapiens* Midasin AAA+ and vWA domains. (A) AlphaFold 3 predicted structure of *H. sapiens* Midasin (residues 1-2500 and 5201-5596 modeled as separate chains), with the N-terminal domain (residues 1-265) light green, six AAA+ domains (residues 266-2420) light brown, beginning of α-helical domain (residues 2421-2500) yellow, and finger-vWA domain (residues 5201-5596) pink. Also included in the prediction were six ATP molecules (shown as sticks) and seven Mg^2+^ ions (shown as gray spheres). (B) View as in panel (A), colored by local confidence (pLDDT). (C) AlphaFold 3 predicted aligned error (PAE) plot for the *H. sapiens* Midasin structure prediction. The location of the finger domain is indicated with an arrow at top.

**Figure S15.**
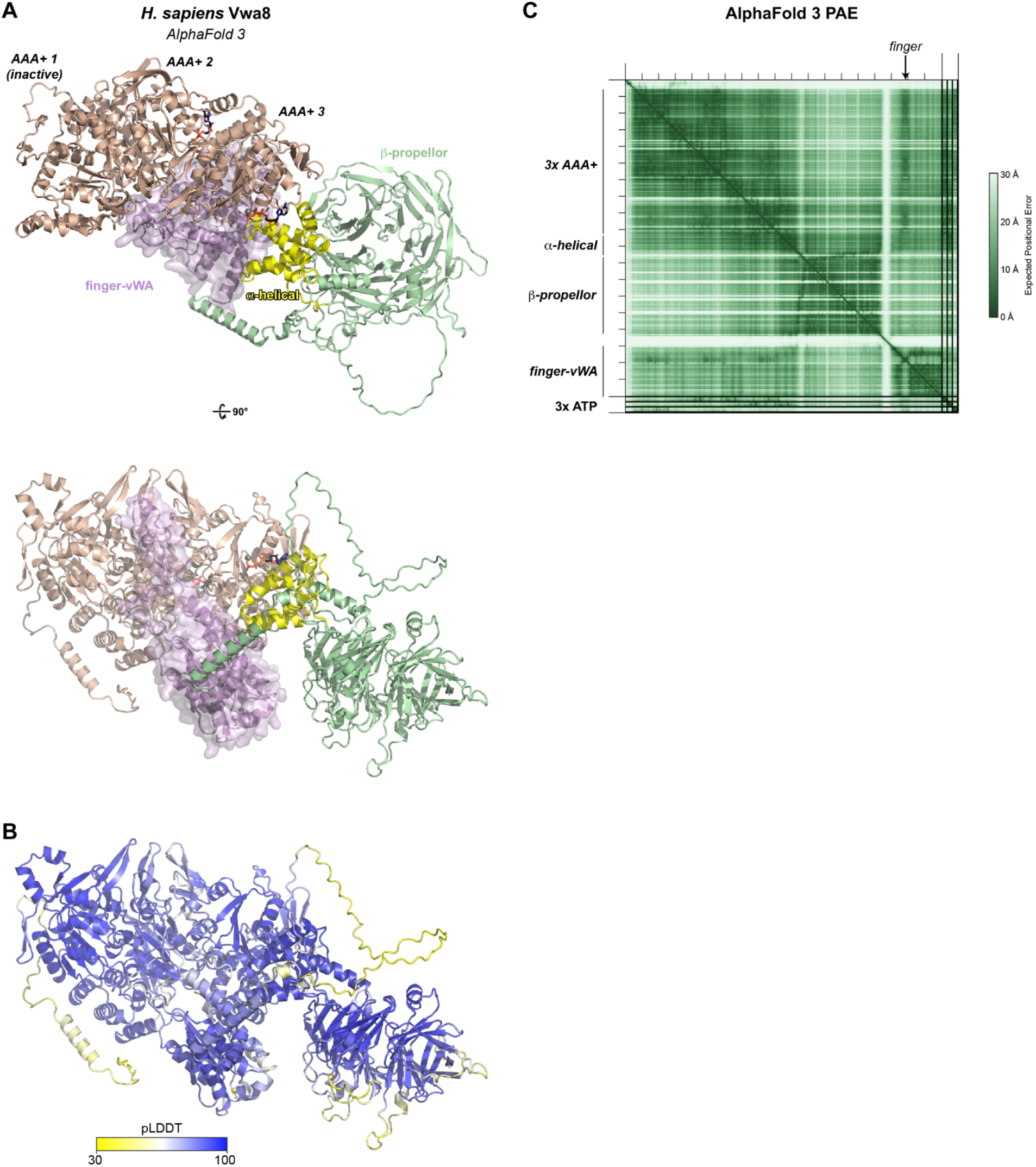
AlphaFold prediction of *H. sapiens* Vwa8. (A) Two views of an AlphaFold 3 predicted structure of *H. sapiens* Vwa8, with three N-terminal AAA+ domains light brown, α-helical domain yellow, β-propeller domain light green, and finger-vWA domain pink. Also included in the prediction were three ATP molecules (shown as sticks) and three Mg^2+^ ions (shown as gray spheres). AlphaFold did not place an ATP·Mg^2+^ in the ATP-binding site of AAA+ domain 1, and close inspection of this site indicates that the Walker A and B motifs are not conserved in this site, likely rendering it inactive. The ATP-binding sites of AAA+ domains 2 and 3 are intact, suggesting that they are active. The three AAA+ domains pack against one another in a manner suggesting that they form a homohexameric assembly, either with a second protomer of Vwa8 or with an unidentified partner protein. (B) View as in panel (A), colored by local confidence (pLDDT). (C) AlphaFold 3 predicted aligned error (PAE) plot for the *H. sapiens* Vwa8 structure prediction. The location of the finger domain is indicated with an arrow at top.

## Supplemental Tables

**Table S1. Genomic neighborhoods of bacterial SpdB homologs**

***See attached Excel workbook***

**Table S2. AlphaFold 2 interaction screening for *E. coli* SpdA, SpdB and SpdC**

*See attached Excel workbook*

**Table S3.**
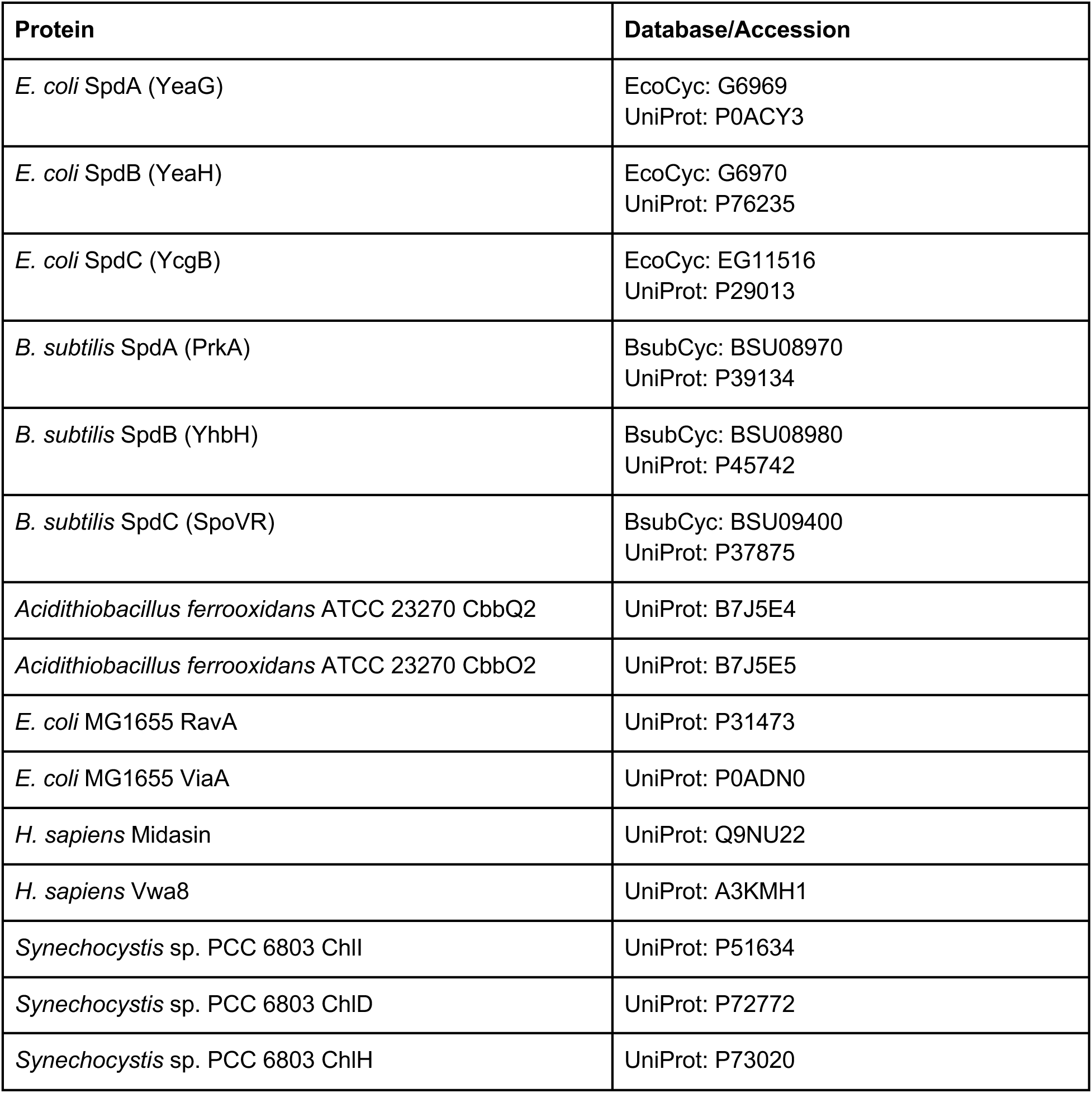
Proteins used in this study.

| **Protein** | **Database/Accession** |
|---|---|
| *E. coli* SpdA (YeaG) | EcoCyc: G6969<br>UniProt: P0ACY3 |
| *E. coli* SpdB (YeaH) | EcoCyc: G6970<br>UniProt: P76235 |
| *E. coli* SpdC (YcgB) | EcoCyc: EG11516<br>UniProt: P29013 |
| *B. subtilis* SpdA (PrkA) | BsubCyc: BSU08970<br>UniProt: P39134 |
| *B. subtilis* SpdB (YhbH) | BsubCyc: BSU08980<br>UniProt: P45742 |
| *B. subtilis* SpdC (SpoVR) | BsubCyc: BSU09400<br>UniProt: P37875 |
| *Acidithiobacillus ferrooxidans* ATCC 23270 CbbQ2 | UniProt: B7J5E4 |
| *Acidithiobacillus ferrooxidans* ATCC 23270 CbbO2 | UniProt: B7J5E5 |
| *E. coli* MG1655 RavA | UniProt: P31473 |
| *E. coli* MG1655 ViaA | UniProt: P0ADN0 |
| *H. sapiens* Midasin | UniProt: Q9NU22 |
| *H. sapiens* Vwa8 | UniProt: A3KMH1 |
| *Synechocystis* sp. PCC 6803 ChlI | UniProt: P51634 |
| *Synechocystis* sp. PCC 6803 ChlD | UniProt: P72772 |
| *Synechocystis* sp. PCC 6803 ChlH | UniProt: P73020 |

**Table S4.** Top DALI PDB25 hits for *E. coli* SpdC AlphaFold 3 structure prediction.

| **PDB ID** | **Chain ID** | **Z score** | **Cα r.m.s.d.** | **Alignment length (AA)** | **Description** |
|---|---|---|---|---|---|
| 3O0Y | A | 11.8 | 3.5 | 186 | Putative lipoprotein |
| 3U24 | A | 11.0 | 3.3 | 175 | Putative lipoprotein |
| 3IUK | A | 10.9 | 3.4 | 186 | M32 carboxypeptidase-like protein |
| 7V9O | A | 10.0 | 5.3 | 181 | Alanine aminopeptidase |
| 1Z1W | A | 9.9 | 5.5 | 176 | Tricorn protease interacting factor F3 |
| 4FGM | A | 9.8 | 6.4 | 192 | Aminopeptidase N family protein |
| 7PFS | A | 9.3 | 12.6 | 198 | Endoplasmic reticulum aminopeptidase 2 |
| 6OIU | C | 9.3 | 6.9 | 191 | Aminopeptidase N |
| 7Q3Y | A | 9.2 | 14.3 | 191 | Angiotensin-converting enzyme |
| 7BB9 | A | 9.1 | 5.1 | 181 | Lugdulysin (zinc metallopeptidase) |

**Table S5.** Strains used in this study. The parent strain for all strains used in this work is *E. coli* K-12 NCM3722 (69) unless otherwise noted. All knockout strains were generated by P1 transduction using phage generated from the *E. coli* BW25113 Keio collection strains (87).

| Name | Genotype | Description | Reference |
|---|---|---|---|
| NCM3722 | Wild type |  | (69) |
| HE845 | Δ*spdA::kan<sup>a</sup>* |  | (10) |
| HE980 | Δ*spdA::frt<sup>b</sup>* pZS-*spdA<sup>+</sup>* |  | This work |
| HE982 | Δ*spdA::frt* pZS-*spdA1* | A strain expressing SpdA mutant carrying E263A | This work |
| HE988 | Δ*spdB::kan* |  | This work |
| HE1127 | Δ*spdB::frt<sup>b</sup>* pZS-*spdB<sup>+</sup>* |  | This work |
| HE1128 | Δ*spdB::frt* pZS-*spdB1* | A strain expressing SpdB mutant carrying D256A S258A S260A | This work |
| HE1115 | Δ*spdC::kan* |  | This work |
| HE1129 | Δ*spdC::frt* pZS-*spdC<sup>+</sup>* |  | This work |
| HE1130 | Δ*spdC::frt* pZS-*spdC1* | A strain expressing SpdC mutant carrying E95A | This work |
| HE1131 | Δ*spdC::frt* pZS-*spdC2* | A strain expressing SpdC mutant carrying H122A H127A | This work |
| HE1132 | Δ*spdC::frt* pZS-*spdC3* | A strain expressing SpdC mutant carrying E283A | This work |
| HE1113 | Δ*clpA::kan* |  | This work |
| HE1119 | Δ*clpA::kan* Δ*spdA::frt* |  | This work |
| HE1114 | Δ*clpX::kan* |  | This work |
| HE1120 | Δ*clpX::kan* Δ*spdA::frt* |  | This work |
| HE1117 | Δ*clpA::frt* Δ*clpX::kan* |  | This work |
| HE1134 | Δ*clpA::frt* Δ*clpX::kan* Δ*spdA::frt* |  | This work |
| HE1017 | Δ*clpP::kan* |  | This work |
| HE1136 | Δ*clpP::kan* Δ*spdA::frt* |  | This work |
| HE997 | Δ*hslV::kan* |  | This work |
| HE1189 | Δ*hslV::kan* Δ*spdA::frt* |  | This work |
| HE1138 | Δ*lon::kan* |  | This work |
| HE1140 | Δ*lon::kan* Δ*spdA::frt* |  | This work |
| HE1187 | ΔΦ(*clpP:clpX:lon*)*::kan* |  | This work |
| HE1186 | ΔΦ(*clpP:clpX:lon*)*::kan* Δ*spdA::frt* |  | This work |
| HE1230 | Δ*lon::frt* pZS-*lon* |  | This work |

<sup>a</sup>Kanamycin-resistance gene (72, 87).
<sup>b</sup>Flip-out scar of kanamycin resistance gene (72).

**Table S6. Proteomics samples**

*See attached Excel workbook*

**Table S7. Proteomics results**

*See attached Excel workbook*

